# A protein interactome for the last eukaryotic common ancestor illuminates the biochemical basis of modern genetic diseases

**DOI:** 10.1101/2024.05.26.595818

**Authors:** Rachael M. Cox, Ophelia Papoulas, Shirlee Shril, Chanjae Lee, Tynan Gardner, Zoya Ansari, Anna M. Battenhouse, Muyoung Lee, Kevin Drew, Claire D. McWhite, David Yang, Janelle C. Leggere, Dannie Durand, Friedhelm Hildebrandt, John B. Wallingford, Edward M. Marcotte

## Abstract

All eukaryotes share a single-celled ancestor from ∼1.5–1.8 billion years ago, the Last Eukaryotic Common Ancestor (LECA). Roughly half of gene families found in modern eukaryotes were already present in LECA, forming molecular systems that continue to influence genetic diseases and traits today. To investigate these systems, we compared genes across 156 organisms to define a core set of protein-coding gene families likely present in LECA, with a quarter remaining uncharacterized. Integrating >26,000 mass spectrometry proteomics analyses from 31 species, we inferred higher-order complexes among these ancient proteins. This reconstructed interactome reveals both established and novel assemblies, offering a biochemical snapshot of LECA’s organization. Finally, by exploring these ancient protein interactions, we found new human gene-disease associations for bone density and congenital birth defects, illustrating the value of ancestral protein networks for modern functional genetics.

## INTRODUCTION

The last eukaryotic common ancestor (LECA), existing ∼1.5-1.8 billion years ago, was the unicellular ancestor of all extant eukaryotes^1,2^. Its molecular makeup offers insights into the ancient innovations underlying modern eukaryotic cellular complexity. As such, understanding the core genetic toolkit of LECA has been a long-standing goal in genetics and evolutionary biology.

Previous phylogenomic analyses agree that LECA was highly complex and contained most hallmarks of the eukaryotic cell^3^. Synthesis of these studies suggests LECA had at least one nucleus^4,5^ with linear chromosomes and centromeres^6^; an interconnected endomembrane system comprised of an endoplasmic reticulum ^7^, Golgi apparatus^8^, vesicle trafficking system^9^, and nuclear envelope^10^; a dynamic actin- and tubulin-based cytoskeleton^11^ including pseudopodia^12^, centrioles^13^, and at least one cilium^14^; distinct degradative vesicles such as lysosomes^15^ and peroxisomes^16^; and mitochondria capable of both aerobic and anaerobic respiration^2,17^.

While there are many partial descriptions of the genetic content of LECA, including systematic reconstructions of many of LECA’s genes^18–22^, there is as yet no integrated picture of LECA’s proteome or how these proteins interact in higher-order assemblies. Such interactome data are valuable because most cellular processes rely on protein assemblies^23^. Thus, an interactome for LECA would provide a richer portrait than genomic reconstructions alone. Moreover, since proteins are the primary drivers of molecular phenotype^24^, protein interaction networks are valuable tools for inferring protein functions and uncovering genotype-to-phenotype relationships of medical and agricultural interest.

Previous efforts to map eukaryotic protein interactomes systematically have primarily concentrated on opisthokonts (*i.e*., animals and fungi)^25–31^ with the exception of a handful of studies in plants (e.g. ^32–34)^ and protists^35,36^. However, comparing protein networks across more divergent species could provide insights not only into evolutionary conservation, but also the divergence of specific biological processes. For example, conserved machinery can be used in different organism-specific contexts to produce distinct organism-specific phenotypes (e.g. the “phenolog hypothesis”^37^).

Here, we reconstruct the LECA protein interaction network and use it to illuminate human genetic disease. We first derived a preliminary estimate of the LECA protein-coding gene set as a basis for interpreting proteomics data. We then integrated ∼26,000 mass spectrometry experiments across 31 eukaryotes spanning ∼1.8 billion years of evolution, including unicellular and multicellular taxa, parasites and free-living species, and motile and non-motile organisms, representing a diverse sampling of eukaryotic cell types, morphologies, lifestyles, and metabolic capacities. The resulting draft map comprises protein interactions present in LECA and widely conserved across the eukaryotic tree of life. Finally, we leveraged this ancient network to predict human gene-disease phenotypes, validated in frogs and mice. Thus, the LECA interactome provides both a framework for understanding eukaryotic origins and a tool for identifying disease-associated proteins.

## RESULTS AND DISCUSSION

### Inferring the gene content of LECA, the last eukaryotic common ancestor

We inferred LECA’s likely gene content using multiple complementary ancestral reconstruction approaches applied to gene families defined by eggNOG orthogroups^38^. As a computationally tractable baseline, we used Dollo parsimony^39^ to model gene duplication and loss across 156 reference proteomes (see **Supplemental Methods** and supporting Zenodo repository), yielding 10,091 candidate ancestral gene families, hereafter referred to as LECA OrthoGroups (OGs) (**Table S1**). To assess robustness—particularly regarding horizontal gene transfer—we evaluated each orthogroup using two additional phylogenetic frameworks that relax Dollo’s single-origin assumption: Wagner parsimony and gene tree–species tree reconciliation. A substantial fraction of Dollo-inferred OGs (6,429; 64%) were independently supported by these approaches (**Figure S1**). We therefore distinguish a “core” LECA OG set supported by multiple methods and restrict all gene- and pathway-specific biological interpretation to this subset. This consensus approach leverages Dollo’s sensitivity (thereby missing fewer families) while mitigating its biases.

While this LECA gene set is a model-dependent approximation for which caveats exist (see **Supplemental Methods** for discussion of limitations), its size (6 to 10K gene families) is consistent both with previous literature estimates^3,19,21^ and with the notion that early eukaryotes were, at minimum, a fusion of at least two prokaryotes’ worth of genes–an archaeal host and a eubacterial mitochondrial endosymbiont. Within the LECA OGs, we recover genes consistent with eukaryotic features inferred from ancestral trait reconstructions, seen by mapping OGs to known functional annotations^38^. **Figure 1** highlights LECA OG functions associated with the nucleus, endoplasmic reticulum, Golgi apparatus, endosomes, digestive vesicles, transport vesicles, secretory vesicles, mitochondria, cilium and an extensive cytoskeleton likely capable of cell projection.

**Figure 1.**
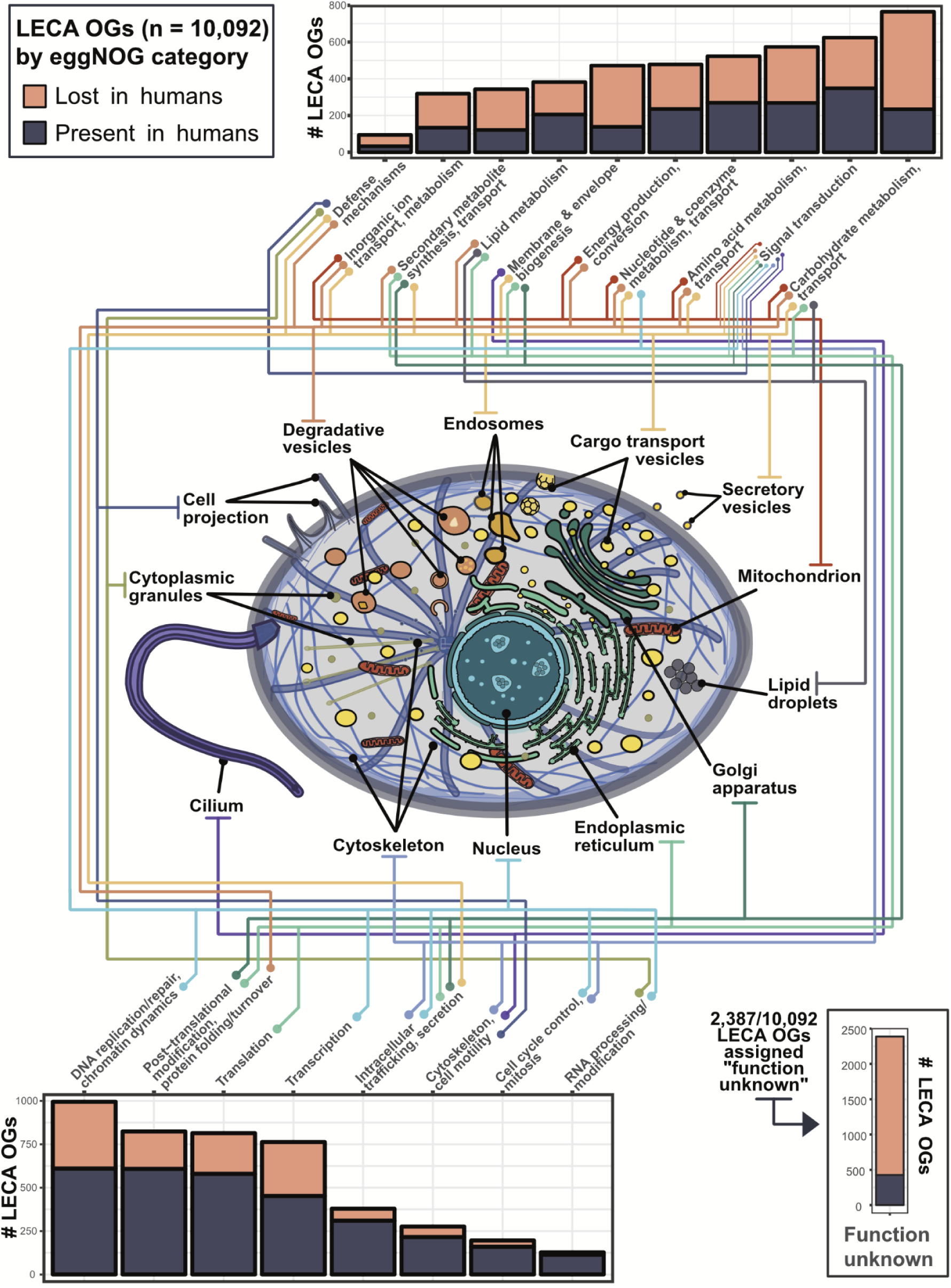
Inferred subcellular organization in LECA, the last eukaryotic common ancestor, based on its estimated gene content. Cell illustration adapted from multiple graphics sourced from SwissBioPics^137^.

Approximately 25% of functionally annotated LECA OGs relate to DNA replication/repair, transcription, translation, and RNA processing, underscoring eukaryote-specific innovations related to the segregation of transcription and translation within the nucleus and cytoplasm, respectively^40^. Examples of these innovations are the spliceosome, nuclear envelope, nuclear pore complex, and transport factors such as karyopherins regulated by the Ran-GTP system^10,41^. Another ∼30% relate to the compartmentalization of energy production, energy conversion, and metabolism^42^. For example, we recover all V-ATPase subunits, a conserved protein complex responsible for the acidification of lysosomes and peroxisomes. Additionally, we traced mitochondrial-specific genes such as TIM and TOM translocases to LECA, in addition to a large suite of mitochondrial carrier family (MLF) proteins and generalist solute carrier (SLC) family proteins. Continuous MLF- and SLC-mediated transport of myriad metabolites across the mitochondrial membrane enables multiple modes of energy conversion^43^, suggesting that LECA had already evolved discriminatory pathways necessary for precise partitioning of varied substrates.

LECA also possessed a rich system of endomembranes, with ∼15% of LECA OGs associated with membrane biogenesis, trafficking, and signaling. These include endosomal coat proteins such as clathrin, adaptin, and the COPI and COPII vesicle coat complexes which facilitate vesicle budding from the Golgi apparatus and endoplasmic reticulum, respectively^9^. Proteins related to signal transduction underwent a massive expansion at the root of eukaryotes, particularly within GTPases, such as the Ras, Ran, Rho, Rab, Arf, and dynamin superfamilies^44^, highlighting the complexity of early signaling pathways and their links to endomembranes. Surprisingly, using eggNOG functional annotations we only found ∼300 LECA OGs (∼3%) associated with the cytoskeleton and cell motility (*i.e.*, a cilium) while previous studies reported ∼500 OGs^45,46^.

Finally, the eggNOG algorithm identified 2,387 LECA OGs (∼25%) of unknown function (**Figure 1, bottom right**), many of which are absent from the human genome. The high proportion of uncharacterized genes and low count of ciliary genes prompted us to attempt to assign function, or at a minimum, subcellular location, to LECA OGs by using annotations for extant proteins in UniProt^47^ (see **Supplemental Methods** and Zenodo repository). This allowed us to recover ∼200 LECA OGs associated with cytoskeletal and cell motility pathways that were previously classified as “function unknown” by the eggNOG algorithm, bringing the total up to the ∼500 expected OGs based on previous literature^45,46^. In sum, we assign tentative UniProt annotations and subcellular compartments to 1,066 of 2,387 LECA OGs originally categorized as “function unknown” by eggNOG.

Overall, slightly more than half of human genes (*n* = 13,571) map to 4,777 unique LECA OGs (of which 93% are core LECA OGs), comparable in scale to previous estimates^48^, and the differences in conservation can be instructive. For example, the human genome appears to have lost a large number of genes associated with carbohydrate metabolism, amino acid metabolism, and membrane biogenesis, including those for isocitrate lyase (ICL) and malate synthase, enzymes of the glyoxylate cycle. Glyoxylate is a highly reactive aldehyde and glyoxylate cycles have been demonstrated in nearly every other branch of life outside of mammals^49,50^. Because of malate synthase loss, human cells must rely on other enzymes to neutralize glyoxylate: alanine-glyoxylate aminotransferase (AGT) in the peroxisome or glyoxylate reductase (GRHPR) in the cytosol. Defects in either human enzyme produce primary hyperoxaluria, *i.e.*, renal disease caused by the failure to detoxify glyoxylate^51^. This example highlights how comparative evolution informs mechanisms of human disease.

### Mapping the LECA protein interactome

We next sought to reconstruct LECA protein organization using co-fractionation mass spectrometry (CFMS), a high-throughput method for measuring protein-protein interactions (PPIs) without tags or affinity reagents (**Figure 2**)^28,52^. CFMS reduces the complexity of a native protein lysate by gently separating protein complexes based on properties such as size, charge, or hydrophobicity, as stably interacting proteins generally co-elute irrespective of the (native) separation. While a single CFMS experiment is insufficient to reliably infer specific protein-protein interactions, integrating observations from orthogonal separations from multiple species, cell types, and tissues confers strong statistical power for inferring conserved interactions^29,31,33^.

**Figure 2.**
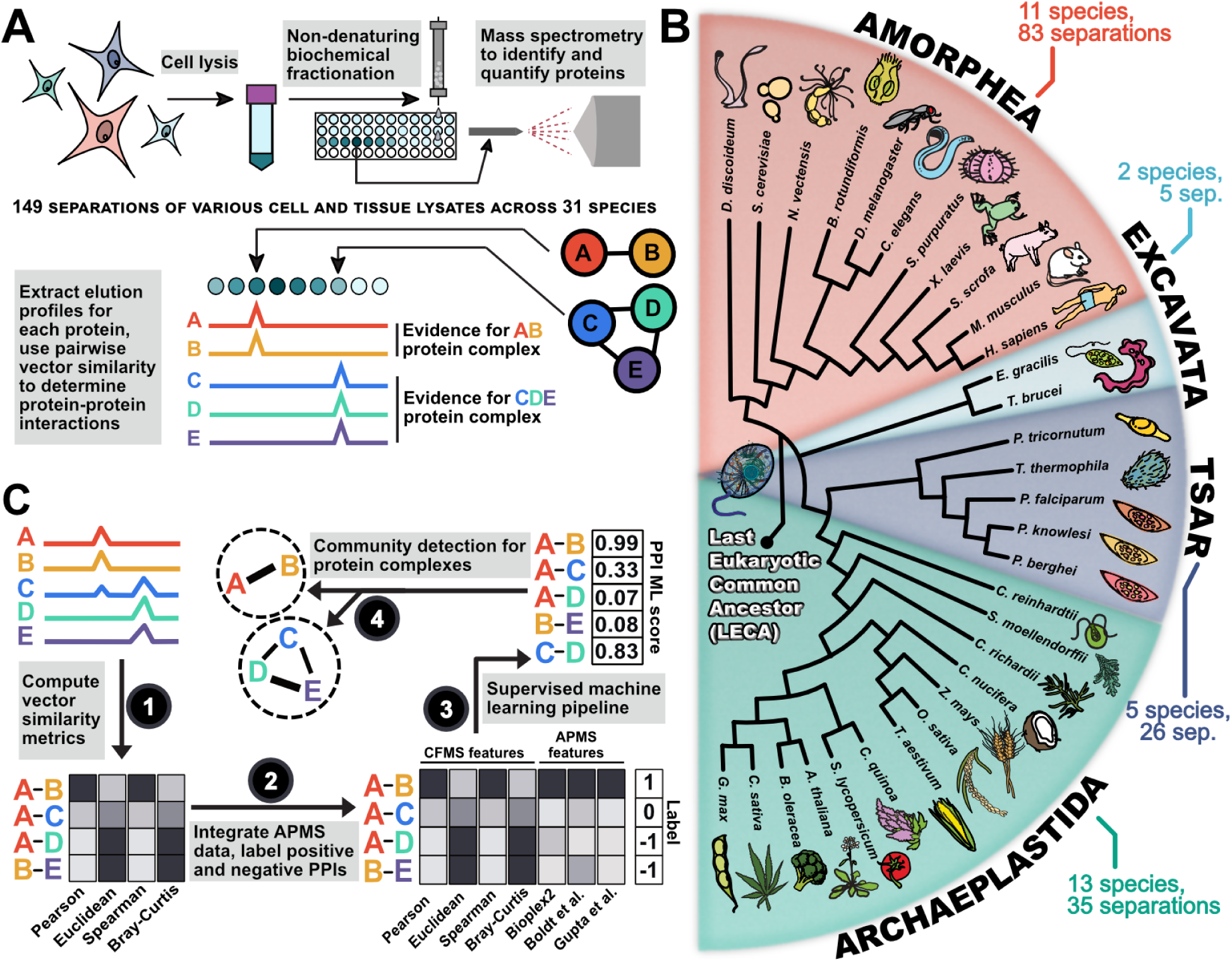
Overview of experimental and computational methods. **(A)** Schematic representation of a co-fractionation mass spectrometry experiment. **(B)** Proteomics data used to construct the LECA interactome included eukaryotes spanning ∼1.8 billion years of evolution. Tree structure is based on ^22^. Branch lengths are not drawn to scale. **(C)** Schematic overview of the approach for computing protein-protein interaction (PPI) features based on CFMS (1) and APMS (2) datasets, scoring conserved PPIs based on these features (3), and clustering scored PPIs into complexes (4).

To detect conserved protein interactions, we integrated raw CFMS data from more than 10,000 individual biochemical fractions^28,29,33,35,36^ across 31 diverse eukaryotic species spanning ∼1.8 billion years of evolution (**Figure 2B**). We also generated new CFMS data for four minimally characterized, phylogenetically diverse unicellular eukaryotes: *Brachionus rotundiformis* (Amorphea, rotifer), *Euglena gracilis* (Excavata, algae), *Phaeodactylum tricornutum* (TSAR, diatom), and *Tetrahymena thermophila* (TSAR, ciliate). Because LECA was ciliated, we expanded our coverage of ciliary proteomes by collecting CFMS data from *Sus scrofa* (Amorphea, pig) tracheal tissue and *Xenopus laevis* (Amorphea, frog) sperm. Experimental details for each separation are provided either in the **Supplemental Methods** section or in the PRIDE database (accessions in the **Resources Table)**.

In all, we measured 379,758,411 peptides that were uniquely assigned to 259,732 orthologous groups (or unique proteins not mapping to orthogroups) across the 31 species, 149 separations, and 10,491 fractions (**Figure 3A**). We augmented our CFMS data with ∼15,000 proteomics experiments^30^ that included affinity purification mass spectrometry (APMS)^53–55^, proximity labeling^56,57^, and RNA-pulldown data^58^. In total, we incorporated data from 26,297 mass spectrometry experiments. We then filtered this data such that we only retained LECA OGs that were strongly observed, in that the sum total of peptide spectral matches (PSMs) across all 149 fractionations was greater than or equal to 150 PSMs. This yielded elution profiles for 5,988 well-measured LECA OGs (∼60% of all LECA OGs and 88% of core OGs).

**Figure 3.**
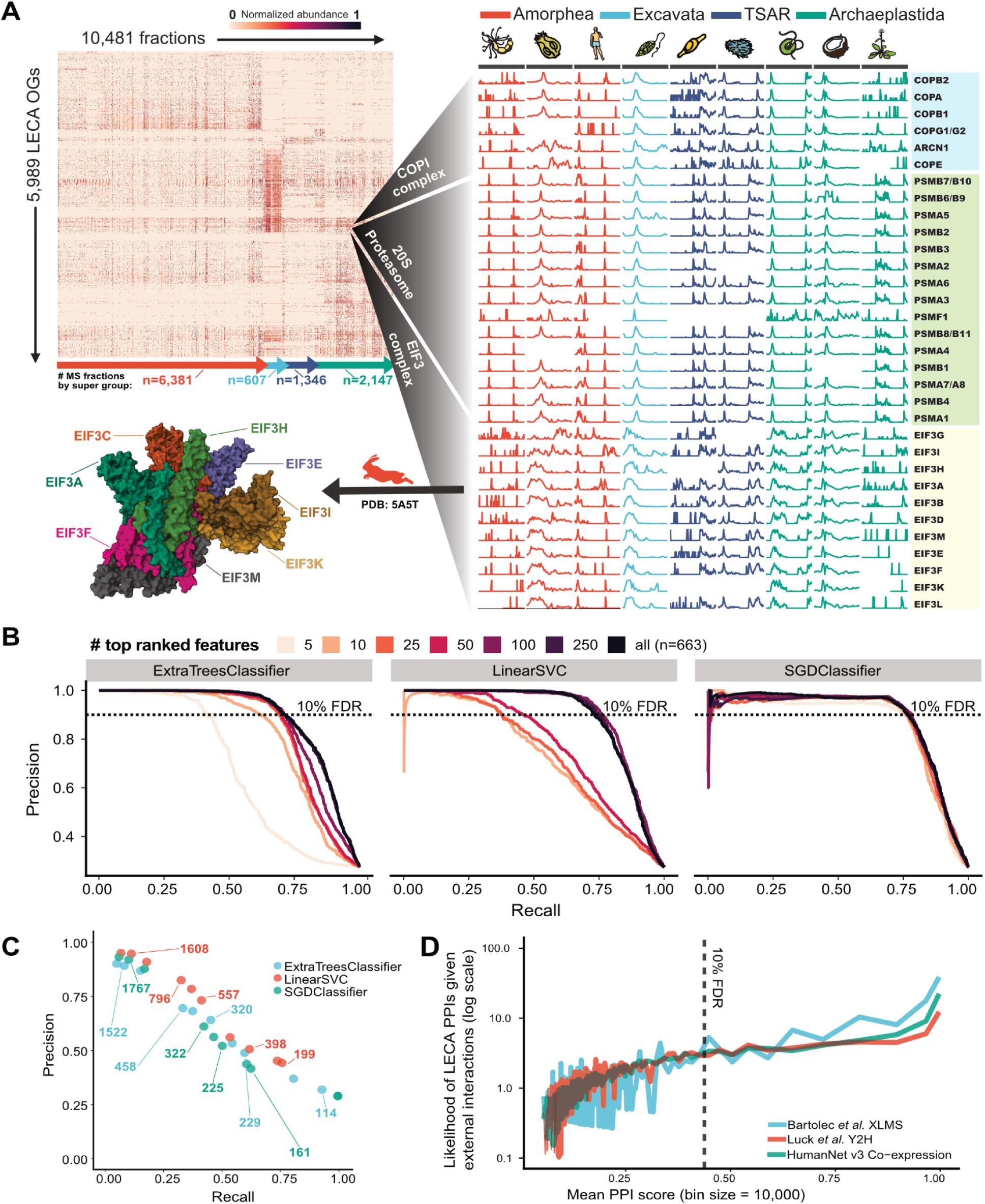
Determining the LECA protein interactome. Co-elution matrix and results of the protein interaction machine learning pipeline. **(A)** Heat map of the filtered elution matrix for 5,989 strongly observed LECA OGs across 10,481 CFMS mass spectrometry fractions (left) and a blow-up of elution vectors for the COPI, 20S proteasome, and eukaryotic initiation factor 3 complexes for a subset of species (right). **(B)** Precision-recall performance of three classifiers trained with increasingly larger sets of ranked features. **(C)** Precision-recall curves for reconstructing known protein complexes defined by a walktrap algorithm, where pairwise PPI scores from each classifier are used as input. Points are labeled with the total number of protein clusters (complexes) constructed at each point in the hierarchy. **(D)** The likelihood of LECA PPIs being present in external protein-protein or mRNA coexpression networks as a function of LECA PPI score, showing agreement increases with scores.

While many complex subunits have co-elution profiles with visually detectable correlation (as for the members of the COPI vesicle coat complex, 20S proteasome, and eukaryotic initiation factor 3 in **Figure 3A, right**), a computational framework is required for systematic identification and to properly control for false positives (**Figure 2C**). To this end, we employed a supervised machine learning pipeline trained on our data for known protein complexes. We assembled a set of 1,499 known complexes from two databases that record PPIs for a variety of eukaryotic species^59–61^ (see **Supplemental Methods**). We then assessed a number of interaction prediction models generated by three different classification algorithms: extremely randomized trees (“ExtraTreesClassifier”), linear support vector (“LinearSVC”), and stochastic gradient descent (“SGDClassifier”).

We constructed models with variable numbers of features derived from the datasets, ranking features on a classifier-by-classifier basis, and evaluated performance by measuring the precision and recall of known complexes withheld from the training data (**Figure 3B**).

Initial models achieved 5-38% recall at 90% precision, similar to previous large-scale protein interaction maps^30,31,33^. However, after implementation of a custom data stratification approach (see **Supplemental Methods**), performance improved to 40-90% recall at 90% precision. We suspect that this large jump in performance stemmed from a combination of the large volume of high quality mass spectrometry data being integrated and the data stratification approach, which we found significantly reduced overfitting (see **Supplemental Methods** for additional discussion of challenges and limitations.)

As the CFMS datasets intrinsically capture higher order multiprotein assemblies (as in **Figure 3A**), we sought to define these assemblies by clustering the LECA OGs based on the measured pairwise interactions. Using each classifier and its best performing feature set, we identified PPIs with a 10% false discovery rate (FDR) threshold (see Zenodo repository) and hierarchically clustered the proteins into complexes on the basis of these interactions (weighted by their confidence scores) using an unsupervised community detection “walktrap” algorithm^62^, which has previously been shown to perform well on CFMS datasets^33^. Rather than rely on a single clustering threshold for determining protein complexes, we defined a hierarchy of protein interactions (**Table S2**), reflecting the natural tendency of proteins to associate into sub-complexes, which assemble into larger complexes, and bind accessory factors. The walktrap procedure also determined an “optimal” number of subcommunities to range between 100-400, differing significantly by classifier. These large clusters capture the general organization of eukaryotic cells (e.g., partitions include a large spliceosomal cluster, a large chromosomal maintenance cluster, clusters broadly associated with cilia, etc).

As a positive control, we noted that this approach successfully delineated known complexes. For example, a large spliceosome cluster was demarcated into LSM, PrP19 complex, and U4/U6 x U5 tri-snRNP complexes with increased granularity (**Table S2**). To quantify walktrap performance, we calculated precision and recall for each cluster at each level of the hierarchy for each classifier (**Figure 3C**), observing a tradeoff where finer subcommunities have increased precision but decreased recall for capturing known complexes. Overall, the support vector classifier (**Figure 3C**, red) netted the highest quality protein complexes and was chosen as our final model. The full hierarchy of LECA protein associations into complexes is reported in **Table S2**.

We assessed how well our final protein interaction model agreed with independent studies that defined interactions using orthogonal approaches. We observe that protein pairs within our highest scoring threshold (≤10% FDR) are significantly more likely than random chance to agree with yeast 2-hybrid (Y2H)^63^, mRNA co-expression^64^, and cross-linking mass spectrometry (XLMS)^65^ interactions (**Figure 3D**), and performed comparably with a previous interaction map in plants^33^. The resulting final LECA complexome consists of the highest confidence 109,466 pairwise interactions between 3,193 unique OGs (of which 98% are core LECA OGs), hierarchically assembled into 199 (less granular) to 2,014 (more granular) protein complexes, which are portrayed schematically in **Figure 4**.

**Figure 4.**
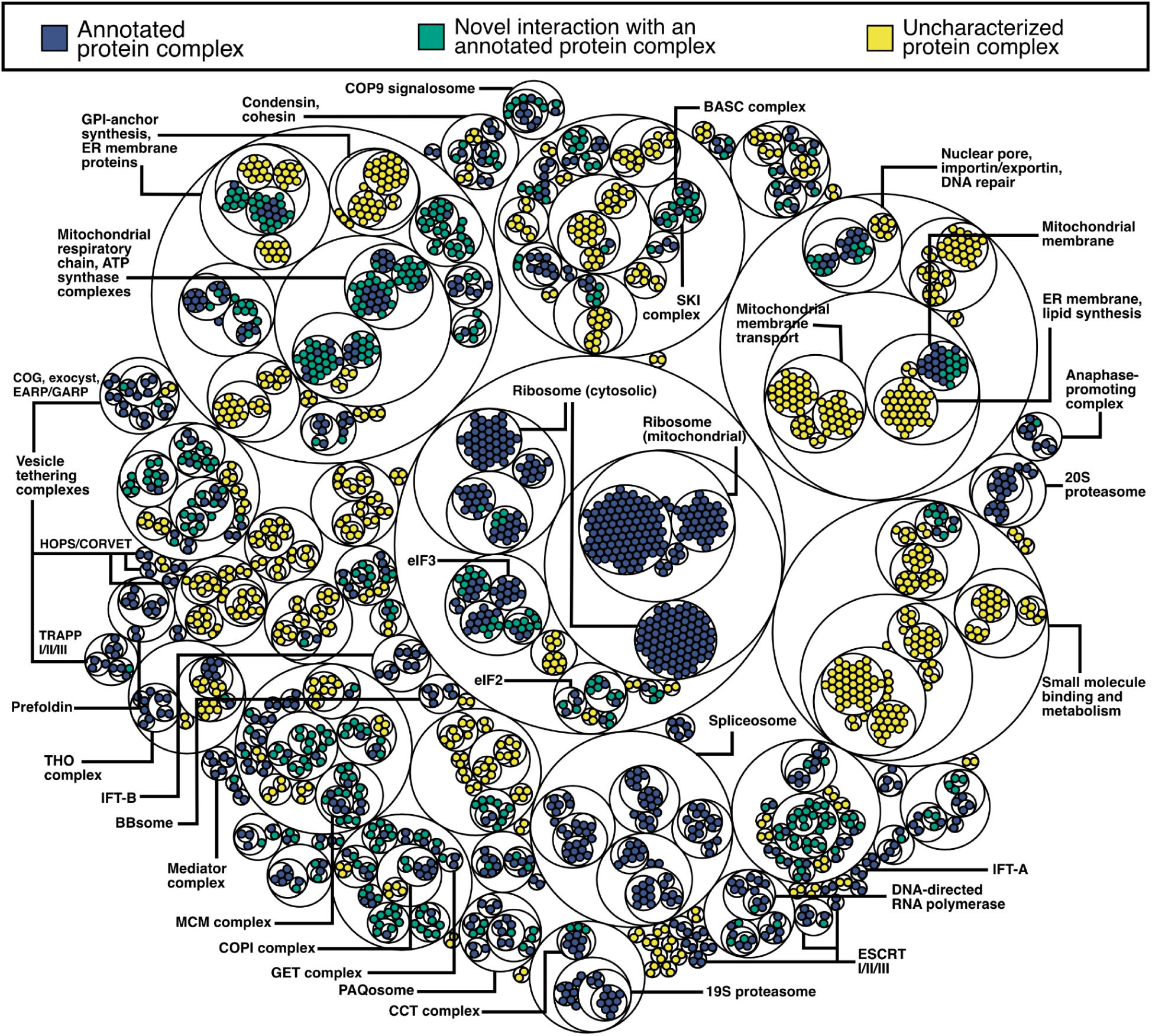
Visualizing hierarchical clustering of protein complexes for a subset of the conserved eukaryotic interactome. The smallest diameter circles correspond to individual proteins, colored by whether the proteins in a cluster are known to interact with each other (blue), whether a novel protein is interacting with a known complex (green), or whether all the associations within a cluster are uncharacterized (yellow).

Importantly for our inference of conserved protein interactions, our method does not rely on any single ancestral reconstruction model. Each interaction in the final network requires independent proteomic evidence from multiple major eukaryotic lineages, such that the interaction map is driven by experimental co-fractionation data rather than by phylogenetic modeling alone. With this experimentally determined set of conserved LECA PPIs in hand, in the next sections, we next examined the extent to which our interactome both recapitulates previously hypothesized and discovers new LECA complexes. Current phylogenomic studies hypothesize myriad protein assemblies at the root of eukaryotes^6,9,10,22,66–68^. We focused on ancient protein assemblies related to intracellular trafficking and cell projection because these are highly relevant to modern human disease^69–71^ and highlight below multiple examples where we uncovered previously undescribed interactions.

### Deep conservation and loss of vesicle tethering complexes

One hallmark of eukaryotic cells is intracellular trafficking by cargo-laden vesicles that bud from one compartment and fuse to another, supported by coat proteins (e.g. clathrin, COPI, and COPII), membrane-anchored SNARE proteins that facilitate fusion^72^, and tethering factors that ensure target specificity^73^. Many compartment-specific tethering modules were likely present in LECA (as in ^66^), including the ER-associated TRAPP-I, TRAPP-II, and TRAPP-III complexes, the Golgi-associated retrograde protein (GARP), the conserved oligomeric Golgi (COG) complexes, the endosome-associated recycling protein (EARP), and the endolysosomal homotypic fusion and protein sorting (HOPS) complexes. However, the extent to which particular protein interactions are conserved remains unknown. In our LECA interactome, we recover all of these core tethering assemblies, along with some unexpected members (**Figure 5A**). These inferences are based on interactions observed across multiple supergroups and restricted to core OGs.

**Figure 5.**
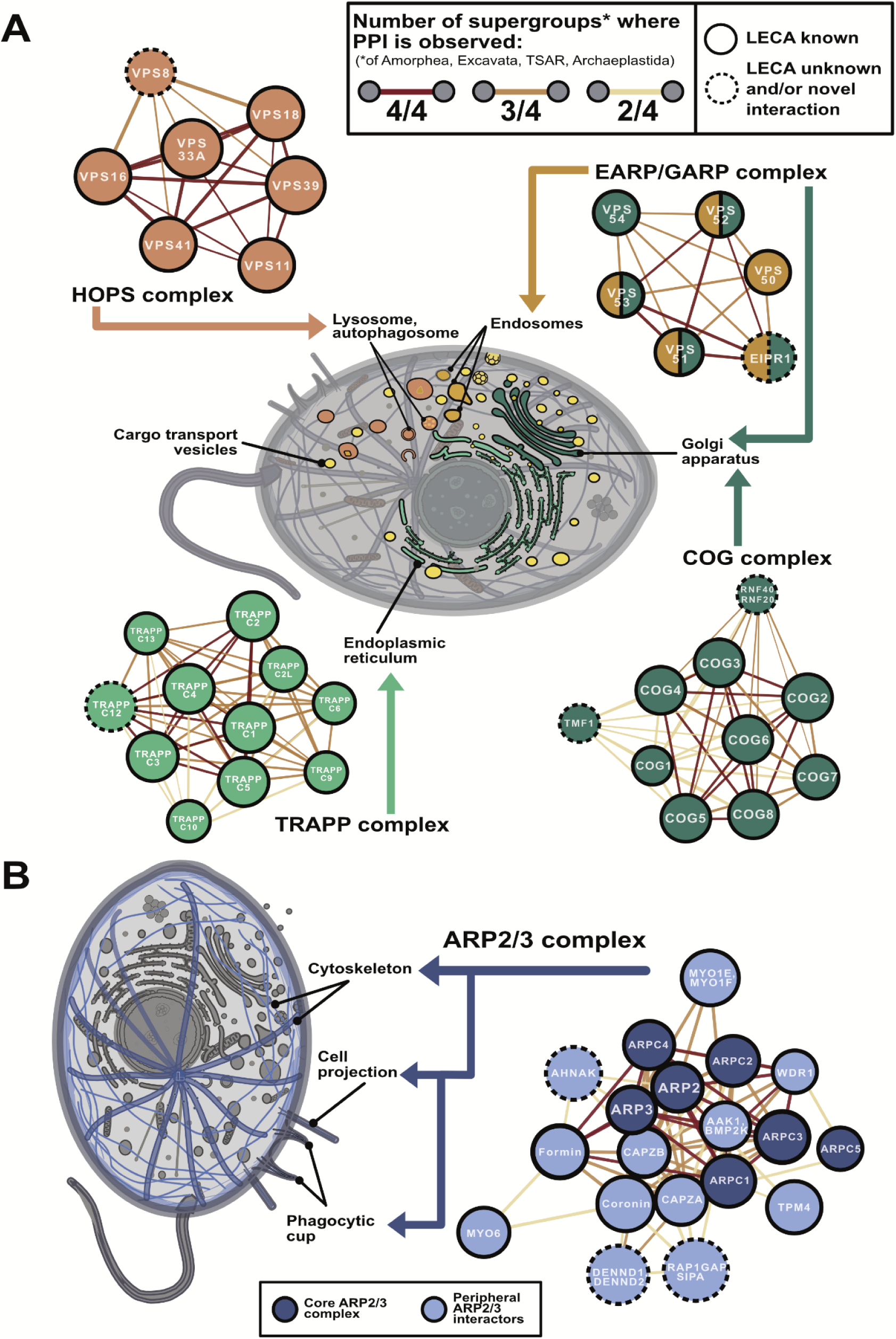
Notable LECA systems related to vesicle tethering and cell projection. **(A)** Node colors for each vesicle tethering complex correspond to their primary subcellular localization: endoplasmic reticulum (light green), Golgi apparatus (dark green), digestive vesicles (orange), or endosomes (yellow). **(B)** Dark and light blue nodes depict core and peripheral cell projection components. In both **(A)** and **(B)**, edges between proteins are colored by the number of eukaryotic supergroups in which the interaction is observed, *i.e.* possesses elution vectors exhibiting a minimum Pearson r of 0.3 within species in two or more of the four eukaryotic supergroups Amorphea, Excavata, TSAR, and/or Archaeplastida (see Figure 2), with darker red edges indicating interactions observed in all supergroups considered.

The GARP and EARP complexes are closely related and share three subunits (VPS51, VPS52, VPS53)^74^. Localization is conferred by additional subunits: VPS50 for endosomes and VPS54 for the Golgi^74^. We observe strongly conserved interactions between all of these subunits, in addition to EIPR1 (EARP and GARP complex-interacting protein 1) (**Figure 5A, top right**). The interaction of EIPR1 with GARP/EARP was only recently discovered, first in high-throughput screens of human proteins^53,55^ and then confirmed in human neuroglioma cells^75^. While EIPR1 is widely conserved, we find that its interaction with GARP/EARP is also ancient and likely traces back to LECA.

In modern eukaryotes, the HOPS complex shares four of its six subunits (VPS11, VPS16, VPS18, VPS33) with the related CORVET complex, while the remaining two (VPS39 and VPS41) are unique to HOPS. In yeast, CORVET subunits direct the fusion of early and recycling endosomes while HOPS directs the fusion of late endosomes, lysosomes, and autophagosomes^76^. We observe conserved interactions between VPS8 (previously thought to be CORVET-specific) and the VPS16, VPS18, VPS39 and VPS41 subunits of the HOPS complex (**Figure 5A, top left**), raising the possibility that a single HOPS-like complex in LECA may have governed endolysosomal vesicle fusion, with subsequent lineage-specific duplication and specialization of subunits for different compartments.

Analogously, the ER/Golgi-associated TRAPP complex is thought to be composed of five core subunits (TRAPPC1-5) with additional subunits in distinct TRAPP-I (TRAPPC6), TRAPP-II (TRAPPC9, TRAPPC10, TRAPPC13), and TRAPP-III (TRAPPC8) complexes. However, the number and identity of proteins in TRAPP-I/II/III vary by species^77^. In our ancient interactome, we observe strong interactions between each member of the core C1-C5 complex, conserved across all sampled eukaryotic supergroups. Unexpectedly, we find pan-eukaryotic evidence for TRAPPC12 in this core complex, previously thought to be metazoan-specific^78,79^. The remaining interactions are differentially lost in specific eukaryotic lineages, with TRAPPC10 absent in all five sampled TSAR species. Our data, combined with conflicting literature on the exact composition of TRAPP-I/II/III, thus suggests an ancient and flexible core complex where subunits differentially specialize along different eukaryotic branches.

The eight-subunit COG assembly governs retrograde intra-Golgi trafficking and comprises two heterotrimeric subcomplexes (COG2-4 and COG5-7) linked by a COG1-COG8 heterodimer^80^. Our LECA interactome recapitulates this assembly and includes interactions with TMF1 and the ubiquitin ligase complex RNF20-RNF40 (**Figure 5A, bottom right**). TMF1-COG interactions have only been previously observed for metazoan COG2 and COG6^81^, but our data show confident TMF1-COG interactions spanning Amorphea and Archaeplastida, with TMF1 lost in Excavata and TSAR. RNF20-RNF40 is generally described in nuclear roles like histone ubiquitination, transcription regulation, and DNA damage repair^82,83^, but has been also linked to the Golgi-associated adapter protein WAC^84^, involved in Golgi membrane fusion^85,86^. We see conservation of RNF20-RNF40 interactions with COG across Amorphea, TSAR, and Archaeplastida, suggesting that nuclear-repurposing of this ubiquitin ligase complex may be a recent mammalian innovation.

Thus, the LECA interactome reveals conservation and specialization of eukaryotic vesicle tethering complexes. We identified unexpected ancient interactions, such as those involving EIPR1 with GARP/EARP, TMF1 and RNF20-RNF40 with COG, and TRAPPC12 with the TRAPP complex, and additionally saw evidence for flexible and lineage-specific adaptations. Given this utility for examining evolutionary conservation and diversification of LECA-associated complexes, we next applied it to shed light on a central question of LECA evolution that is in dispute.

### Primordial origins of cell projection and phagocytosis

While phagocytosis is widely observed across eukaryotes, it remains debated whether LECA could recognize and engulf large particles. Contention stems from how the first eukaryotic common ancestor (FECA) acquired the alpha-proteobacterial precursor of the mitochondrion, *i.e.*, whether FECA was akin to a phagocytosing archaeon or a more “simple” prokaryote that existed in protracted syntrophy with an alpha-proteobacterium^87–89^. Existing phylogenomic investigations into the origins of phagocytosis are conflicting; at least three independent studies conclude phagocytosis probably evolved independently in multiple eukaryotic lineages^90–92^, while others argue that the trait was present in LECA and the absence of phagocytosis in certain eukaryotic groups is due to secondary loss when adapting to new niches^93,94^. Recent investigations into Asgard archaea, the sister group to eukaryotes, reveal a dynamic actin-based cytoskeleton composed of F-actin assemblies, actin-related proteins (Arps), and actin-binding proteins such as profilins and gelsolins capable of modulating eukaryotic actin^95–97^, suggesting that FECA may have had phagocytic capacity.

Within our LECA gene set, we find an extensive complement of OGs associated with both cell projections (pseudopodia, lamellipodia, filopodia) and phagocytosis (phagocytic cups and phagosomes) (**Figures 1, 5B**). Specifically, LECA appears to have had Rho GTPases such as RAC/CDC42, formins, coronins, cofilins, gelsolins, and proteins associated the ARP2/3, ENA/VASP, WASP, SCAR/WAVE, and PI3K complexes. For example, the LECA interactome includes all seven subunits of the ARP2/3 complex, responsible for the cytoskeletal rearrangement required for cell projection, clustering with protein complexes involved in cell protrusion and proteins critical for phagocytosis in extant eukaryotic cells (**Figure 5B**). The core ARP2/3 complex consists of the proteins ARP2, ARP3, ARPC1, ARPC2, ARPC3, ARPC4, and ARPC5. Interestingly, in our data, ARPC5 is the most peripherally associated component in the cluster and has been completely lost in all species sampled within Excavata and TSAR.

Combined with previous evidence^98–100^, these genes suggest that the ancestral eukaryotic cell was likely capable of pseudopod formation and projection-based motility, despite the lack of UniProt annotations in species outside Amorphea (**Figure S2**). However, because cell projections and phagocytosis share underlying molecular machinery, it is less clear if the presence of these systems necessarily imply a phagocytosing LECA, and more evidence is required to conclude that phagocytosis is ancestral at the root of eukaryotes. To address this, we explored the conservation of additional interactions.

Among peripheral interactors of the ARP2/3 complex, we observe CAPZA and CAPZB forming the heterodimeric F-actin capping complex, an essential regulator of actin nucleation that restricts elongation^101^, as well as formins and coronins known to promote elongation^98,102^. We also find interactions with WDR1, a promoter of cofilin-mediated actin severing^103^ that assists both actin polymerization and depolymerization^104^. Research strongly implicates these systems in Amorphean phagocytosis: Coronins are strongly enriched in phagocytic cups and defects result in impaired phagocytosis in both *Dictyostelium* and mammalian cells^105–107^. Furthermore, we observe interaction evidence in at least two major eukaryotic supergroups consistent with the reported roles of AAK1 in receptor mediated endocytosis^108^ and unconventional myosin in phagocytosis^109–112^. Taken together, our results strongly support a phagocytosing LECA.

### The LECA interactome reveals a ciliary mechanism for *EFHC2*-associated renal failure

Conservation of protein interactions over billions of years implies strong constraint and likely pathogenicity when disrupted. Thus, studying genetic variation through the lens of conserved protein interactions should clarify mechanisms of genetic disease development, tolerance, and resilience. We therefore expect LECA protein interactions to offer direct insights into human disease genetics, and by similar logic, into genotype-phenotype relationships of other eukaryotes.

Slightly more than half of human genes date back to LECA. Where these conserved genes have been characterized, they are responsible for a large and diverse subset of major human diseases, spanning developmental disorders, cancers, chronic respiratory diseases, neurodegenerative conditions, and motor disorders (**Figure 6A**). For example, of the ∼100 human genes linked with deafness, nearly ¾ were present in LECA (**Figure 6A**). While some human diseases are “new”, evolutionarily-speaking, such as deficiency of the animal-specific pituitary hormone, many other diseases, such as ciliary dyskinesia, arise nearly entirely from genes in LECA OGs. In order to test the utility of our data for illuminating the biology of extant species, we next asked if the LECA interactome could be leveraged to predict human disease mechanisms and novel gene-disease relationships.

**Figure 6.**
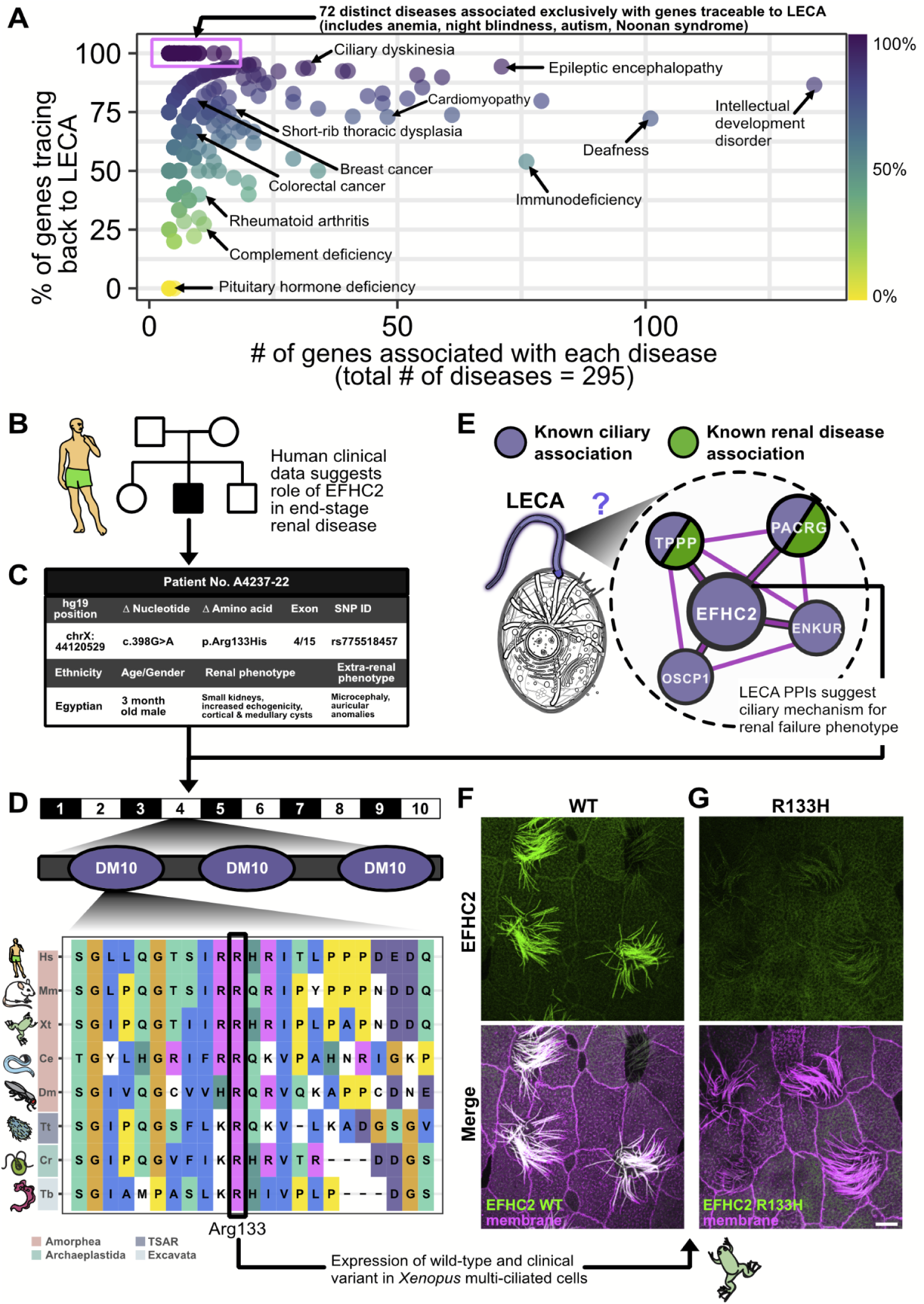
LECA protein interactions suggest mechanisms of genetic disease, as for end stage renal disease gene *EFHC2*, identified by whole exome sequencing and confirmed to have a ciliary etiology. **(A)** Causal genes for human diseases are frequently ancient, as shown by plotting gene-disease relationships from OMIM, with each point representing a unique disease group with an associated number of genes (x-axis) and age, determined as the percentage of genes in LECA OGs (y-axis). **(B)** Pedigree of the index family A4237. Squares represent males, circles females, black shading the affected proband individual A4237-22 included in whole-exome sequencing (WES), and white shading the unaffected parents and siblings. **(C)** Summary of the phenotype and recessive disease-causing R133H *EFHC2* variant identified by WES. **(D)** Location of Arginine 133 in relation to *EFHC2* exon/intron (black/white) structure and DM10 protein domains (purple), and its deep evolutionary conservation. **(E)** EFHC2-containing ciliary complex uncovered in the LECA interactome. **(F)** Localization of GFP-EFHC2 to axonemes in *Xenopus* motile cilia. **(G)** Introduction of the R133H mutation results in loss of ciliary localization of GFP-EFHC2, confirmed by co-labeling with membrane-RFP. Scale bar = 10 µm

Approximately 500 LECA OGs are related to cilia, and among the most common ciliopathies are diseases of the kidney^113^. We identified a male infant with microcephaly, seizures, polycystic kidney disease, and end-stage renal failure, and whole exome sequencing and pedigree analyses revealed a significant hemizygous, X-linked G>A variant in *EFHC2* (rs34729789, 11:44148852:G:A) (**Figures 6B-D, S3**). Little is known of the function of *EFHC2*, though it and its paralog *EFHC1* (both belong to the same OG) encode proteins thought to be microtubule inner proteins (MIPs) that function specifically in motile cilia; loss of their orthologues in *Chlamydomonas* or *Tetrahymena* leads to defective ciliary axonemes and/or ciliary beating, but do not disrupt ciliogenesis^114,115^. Because motile cilia are absent from mammalian kidneys, the link between *EFHC2* and this patient’s disease was surprising. We therefore examined our LECA interactome for insights.

We first noted that the patient’s missense variant altered an arginine residue at position 133 to histidine, and this residue is conserved across Archaeplastida, Excavata, TSAR, and Amorphea (**Figure 6D**). Moreover, EFHC2 was closely and exclusively linked in our LECA interactome to other proteins involved in cilia motility (**Figure 6E**). We therefore examined the protein’s localization in *Xenopus* multiciliated cells, and found it to be very strongly localized to ciliary axonemes. By contrast, the disease-associated R133H variant failed to localize to cilia (**Figure 6F, G**), suggesting that defective ciliary localization of this protein contributed to ciliopathic kidney disease in the affected child.

This result prompted us to ask if other cluster proteins, also thought to function in motile cilia, might be implicated in kidney disease. Indeed, previous genomic analyses link both *PACRG*^116^ and *TPPP*^117^ to chronic kidney disease, with *PACRG* specifically linked to end-stage renal disease^116^. This combination of clinical data, the LECA interactome, and specific hypothesis testing in a vertebrate model organism thus links EFHC2 ciliary function to an end-stage renal disease for which the molecular etiology was previously unknown and underscores the power of our comparative evolution strategy.

### Network propagation for systematic ranking of gene-disease relationships

We next sought to score potential disease-causative proteins systematically within our conserved interactome on a disease-by-disease basis. To this end, we used cross-validated network guilt-by-association^118^ to predict novel gene-disease pairs for 109 diseases based on clinically-validated genotype-to-phenotype relationships sourced from the OMIM database^119^ (**Figure 7A**; see **Supplemental Methods**). We measured performance as areas under receiver operating characteristic curves (AUROC), and compared the predictive performance of known disease-associated gene sets versus that of random gene sets. As expected, random predictions yield AUROC scores around 0.5 (**Figure 7A, yellow**). In contrast, LECA-interactome predictions skewed to higher scores (**Figure 7A, blue**), and using a score threshold of 0.7, we made strong new disease candidate predictions for almost one-third of the diseases considered (∼35 Mendelian disorders). As the LECA interactome should also predict phenotypes in other eukaryotes, we performed similar tests for yeast and *Chlamydomonas* mutant phenotypes and observed similarly strong performance (**Figure S4**). Notably, because network propagation operates on experimentally supported conserved interactions, rather than on phylogenetic reconstruction alone, these phenotype predictions are robust to uncertainty in the precise evolutionary origin of individual gene families. Below, we discuss the prediction and validation in animals of two such novel protein associations.

**Figure 7.**
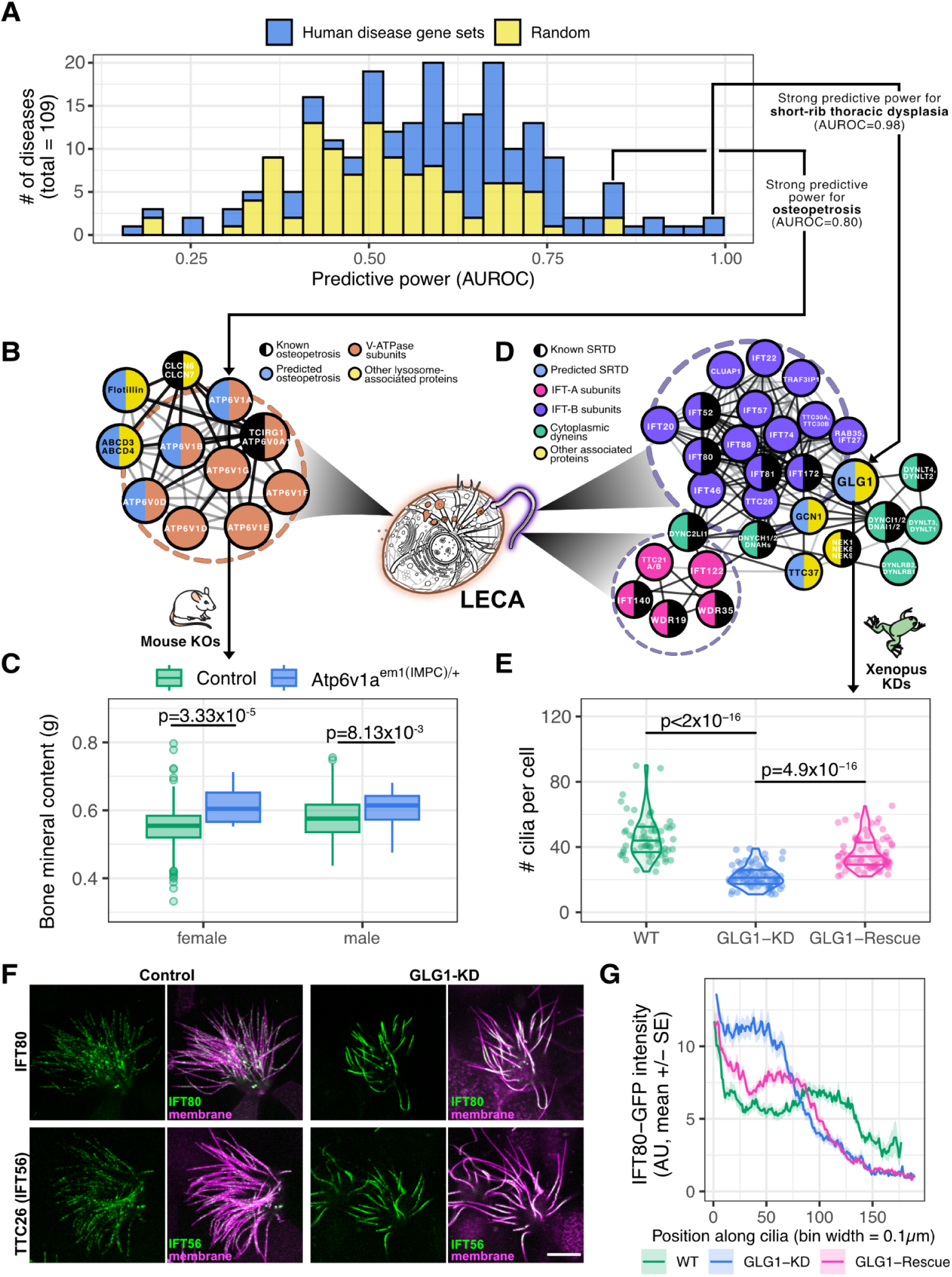
Guilt-by-association in the LECA interactome identifies *ATP6V1A* as causative for osteopetrosis and *GLG1* for short-rib thoracic dysplasia (SRTD). **(A)** Guilt-by-association in the LECA PPI network correctly associates genes to human diseases for roughly ⅓ of the 109 diseases tested, measured as the areas under receiver operating characteristic curves (AUROCs) of leave-one-out cross-validated predictions of known disease genes (light blue) versus random associations (yellow). **(B)** PPI network of genes clinically linked to osteopetrosis (black half-discs; 3 additional genes lie outside this cluster), the highest-ranking new candidates (purple), and their interactions with other V-ATPase subunits that were not indicated for osteopetrosis (orange). **(C)** For the top-scoring gene *ATP6V1A*, the bone mineral content is plotted for knockout (KO) mice with a heterozygous exon deletion in *ATP6V1A* (*n*=8 for each sex, *n*=16 total) compared to healthy control mice (female *n*=834, male *n*=780). Null mice show significantly increased bone density, consistent with the clinical manifestation of osteopetrosis. **(D)** The PPI network of genes clinically linked to SRTD (black half-discs) implicates *GLG1* (yellow) and suggests a ciliary role, based on interactions with intraflagellar trafficking IFT-A (blue) and IFT-B (purple) complexes, cytoplasmic dyneins and dynactins (green), and other interactors (gray). **(E)** Morpholino knockdown (KD) of *GLG1* significantly reduced the number of cilia in *X. laevis* multi-ciliated cells (Bonferroni adjusted t-test *p* < 10^-16^, *n* = 60 control cells, 79 knockdown cells, and 76 rescue cells, 9 embryos per condition over 3 injection replicates) compared to uninjected control animals; rescue by co-injection with a non-targeted *GLG1* allele confirmed specificity. **(F)** In control *Xenopus* multi-ciliated cells, IFT56-GFP and IFT80-GFP, two subunits of IFT-B, are distributed as particles along the ciliary axonemes. However, MO knockdown of *GLG1* leads to the accumulation of IFT-B proteins in the proximal region of axonemes. Scale bar = 10 µm. **(G)** This effect is quantified for IFT80-GFP for 3 cilia per cell for all cells analyzed in panel **(E)**.

### Identification and validation of *ATP6V1A* as a novel candidate for osteopetrosis

Our LECA network propagation approach implicated several vacuolar-type H^+^-ATPase (V-ATPase) proteins in the molecular etiology of osteopetrosis (AUROC ∼0.8), a disorder in which bones grow abnormally and become overly dense^120^ (**Figure 7B**). Given the role of V-ATPases in bone homeostasis by acidifying the space between osteoclasts and bone to help dissolve hydroxyapatite^121^, one might assume that disrupting many V-ATPase subunits increases bone density. However, only three subunits have been implicated in osteopetrosis in humans^119,122^ or mice^123^. The remaining subunits instead display a remarkably broad spectrum of disease associations, including cutis laxa (loose skin) and renal tubular acidosis^124^, neurodegenerative disease, deafness^125^, Zimmermann-Laband syndrome^126^, and even osteoporosis (bone loss)^127^, highlighting the need to elucidate the discrete molecular functions of specific V-ATPase subunits.

Our LECA network propagation approach gratifyingly made precise predictions, linking three specific subunits (ATP6V1A, ATP6V1B, ATP6V0D) to osteopetrosis (**Figure 7B**). To confirm these predictions we examined heterozygous CRISPR-Cas9 knockouts of *ATP6V1A* (performed by the KOMP2 high-throughput mouse phenotyping site at the Baylor College of Medicine, see **Supplemental Methods**) and found these mice showed significantly increased bone mineral content (**Figure 7C**). The effect size was much stronger in female mice (*p* = 0.00003) than in male mice (*p* = 0.00813), echoing previous observations of sexual dimorphism in the body composition of mammals^128,129^. Despite the obvious lack of bones in the single celled last eukaryotic ancestor, then, our examination of the protein interaction network of that organism nonetheless identified a specific mammalian phenotype with one specific subunit from among a large repertoire of closely related genes.

### Ancient interactions suggest new candidate genes for a lethal human ciliopathy

Our highest scoring disease association (AUROC ∼0.98) involved short-rib thoracic dysplasia (SRTD), a severe human ciliopathy characterized by skeletal abnormalities including dysplasia of the axial skeleton that often lethally impairs respiratory function^130^. The disease is strongly associated with proteins in Intraflagellar Transport (IFT), the system moving cargoes into and out of cilia, and this was reflected in our LECA interactome (**Figure 7D**). Our highest scoring non-IFT protein prediction, however, was the Golgi protein GLG1^131^. This was notable because mouse GLG1 mutants display rib defects similar to SRTD^132^, yet the protein has never been implicated in any aspect of ciliary biology.

We therefore explored the function of GLG1 in *Xenopus* multiciliated cells (MCCs), and found it predominantly localized to the Golgi in MCCs, as expected. We observed no localization at basal bodies or in cilia (not shown). Nonetheless, GLG1 knockdown resulted in significant loss of cilia from MCCs, an effect that was specific since it could be rescued by expression of GLG1-FLAG (**Figure 7E**). To ask if this ciliogenesis defect was related to IFT, we performed live imaging of GFP fusions to two IFT proteins. In normal cells, both markers labeled small punctae in axonemes of *Xenopus* MCCs, consistent with previous imaging of IFT in these cells^133,134^. By contrast, GLG1 knockdown cells displayed large accumulations of IFT proteins within axonemes (**Figure 7F, 7G**) that resemble those seen previously after disruption of IFT^133,134^.

Thus, analysis of the LECA interactome made a single, specific prediction of ciliary function for just one among the large array of Golgi-resident proteins, and that prediction was validated by experiments in *Xenopus*. These data provided new insights into the still obscure link between IFT and the Golgi^135,136^ and, moreover, identified a plausible candidate gene for SRTD.

### Conclusions

Conserved ancient protein interactions provide insight into the genetic basis of modern traits and diseases. In this work, we reconstructed macromolecular assemblies of ancient proteins that as yet have only been sparsely described. We defined a core set of likely LECA orthogroups, finding that slightly more than half of human genes can be traced back to this set. We integrated those data with >26,000 mass spectrometry proteomics experiments, capturing hundreds of millions of peptide measurements for hundreds of thousands of proteins in species spanning the eukaryotic tree. Using these data, we reconstructed a high-quality conserved LECA protein interactome. Improved probabilistic gain–loss and DTL models may refine membership at the margins of the LECA gene set, but are unlikely to alter the major conserved complexes defined here, which are supported by direct proteomic evidence across multiple eukaryotic supergroups. This interaction network has formed the core of eukaryotic biology for nearly two billion years, and the dataset gives insights into known protein complexes and novel assemblies.

Consistent with our central premise that the most highly-conserved protein assemblies will tend to be most critical for cell and organism function, the LECA interactome successfully predicts mechanisms of human disease and new gene-disease relationships. We presented evidence for a ciliary mechanism in human *EFHC2*-associated renal failure, identified the V-ATPase subunit ATP6V1A in the etiology of mammalian osteopetrosis, and demonstrated a role for the Golgi protein GLG1 in trafficking IFT-A proteins into cilia as a molecular mechanism for short-rib thoracic dysplasia. Given the richness of these datasets, we expect this approach to extend to traits and diseases in most eukaryotes, while providing insights into specific mechanisms involved due to being anchored in deeply conserved protein activities.

### Limitations of the study

Our reconstruction of LECA’s proteome and interactome has several limitations, including assumptions of the models used to infer ancestral genes, uneven taxonomic and experimental sampling, limited resolution of paralogs, and biases in CFMS favoring abundant, stable complexes. A discussion of these limitations is provided in the **Supplemental Methods**.

## RESOURCE AVAILABILITY

### Lead contact

Requests for further information and resources should be directed to and will be fulfilled by the lead contact, Edward M. Marcotte.

### Materials availability

This study did not generate new materials.

### Data and code availability

All raw and interpreted mass spectrometry data were deposited to the ProteomeXchange *via* the MassIVE repository under the identifiers in the **Resources Table**. Project data files are available at Zenodo (https://doi.org/10.5281/zenodo.10961422). Custom R, Python, Bash, and Perl scripts for analyses and figure generation are available at https://github.com/marcottelab/leca-proteomics and https://github.com/marcottelab/leca_notung_reconciliation.

## ACKNOWLEDGEMENTS

The authors gratefully acknowledge support from the Tetrahymena stock center (Cornell University & Washington University in St. Louis), the Texas Advanced Computing Center (UT Austin) for computing resources, the Knockout Mouse Program (KOMP2) at the Baylor College of Medicine for *Atp6v1a* mice data, Johann Eberhart for rotifer cultures, the UTEX Culture Collection of Algae for algal samples, Maureen Stolzer for assistance with reconciliation, and Angel Syrett and Elinor Marcotte for species illustrations. Research was funded by grants from the National Institute of General Medical Sciences (R35GM122480 to E.M.M. and F31GM143881 to R.M.C), National Institute of Diabetes and Digestive and Kidney Diseases (R01DK068306 to F.H.), National Institute of Child Health and Human Development (R00HD092613 to K.D. and R01HD085901 to J.B.W. and E.M.M.), Army Research Office (W911NF-12-1-0390 to E.M.M.), and Welch Foundation (F-1515 to E.M.M.).

## AUTHOR CONTRIBUTIONS

Design and co-supervision: R.M.C., J.B.W., and E.M.M.

Proteomics experiments: O.P., aided by K.D. and J.C.L.

Data analysis: R.M.C., aided by T.G., Z.A., A.M.B., M.L., K.D., C.D.M., D.Y., and D.D., and guided by E.M.M.

Clinical genetics: S.S. and F.H.

*Xenopus* experiments: C.L., T.G., and J.B.W.

Manuscript initial draft: R.M.C., O.P., J.B.W., and E.M.M.

All authors discussed results and contributed edits.

## SUPPLEMENTAL METHODS

### RESOURCES TABLE

| REAGENT OR RESOURCE | SOURCE | IDENTIFIER |
| --- | --- | --- |
| **BIOLOGICAL SAMPLES** |  |  |
| *Euglena gracilis* | UTEX Culture Collection of Algae | UTEX 753 |
| *Tetrahymena thermophila* | Tetrahymena Stock Center | SB175 (SD01508) |
| Pig trachea (*Sus scrofa*) | Sierra for Medical Science | N/A |
| Diatom (*Phaeodactylum tricornutum*) | UTEX Culture Collection of Algae | UTEX 646 |
| Rotifer (*Brachionus rotundiformis*) | Eberhart lab, UT Austin | S-type |
| **MASS SPECTROMETRY DATA** |  |  |
| *Euglena gracilis* | This work | **PRIDE:** PXD050669 |
| *Tetrahymena thermophila* | This work | **PRIDE:** PXD050671, PXD050672, PXD050674 |
| Pig (*Sus scrofa*) | This work | **PRIDE:** PXD041980 |
| Diatom (*Phaeodactylum tricornutum*) | This work | **PRIDE:** PXD050670 |
| Rotifer (*Brachionus rotundiformis*) | This work | **PRIDE:** PXD050673 |
| *Arabidopsis thaliana* | McWhite et al., 2020 | **PRIDE:** PXD013264, PXD013321, PXD014617 |
| Broccoli (*Brassica oleracea*) | McWhite et al., 2020 | **PRIDE:** PXD013281, PXD013322, PXD013282 |
| *Chlamydomonas reinhardtii* | McWhite et al., 2020 | **PRIDE:** PXD013369, PXD013735 |
| Coconut (*Cocos nucifera*) | McWhite et al., 2020 | **PRIDE:** PXD012865 |
| Fern (*Ceratopteris richardii*) | McWhite et al., 2020 | **PRIDE:** PXD013320 |
| Hemp (*Cannabis sativa*) | McWhite et al., 2020 | **PRIDE:** PXD012969 |
| Maize (*Zea mays*) | McWhite et al., 2020 | **PRIDE:** PXD012810 |
| Quinoa (*Chenopodium quinoa*) | McWhite et al., 2020 | **PRIDE:** PXD013080 |
| Rice (*Oryza sativa*) | McWhite et al., 2020 | **PRIDE:** PXD013213 |
| *Selaginella moellendorffii* | McWhite et al., 2020 | **PRIDE:** PXD013093 |
| Soy (*Glycine max*) | McWhite et al., 2020 | **PRIDE:** PXD013198, PXD013704 |
| Tomato (*Solanum lycopersicum*) | McWhite et al., 2020 | **PRIDE:** PXD013004 |
| Wheat (*Triticum aestivum*) | McWhite et al., 2020 | **PRIDE:** PXD013214, PXD013280, PXD013300 |
| *Plasmodium berghei* | Hillier et al., 2019 | **PRIDE:** PXD009039 |
| *Plasmodium falciparum* | Hillier et al., 2019 | **PRIDE:** PXD009039 |
| *Plasmodium knowlesi* | Hillier et al., 2019 | **PRIDE:** PXD009039 |
| *Trypanosoma brucei* | Crozier et al., 2017 | **PRIDE:** PXD005968 |
| Human (*Homo sapiens*) | Wan et al., 2015 | **PRIDE:** PXD002322, PXD002328 |
| Worm (*Caenorhabditis elegans*) | Wan et al., 2015 | **PRIDE:** PXD002319 |
| Slime mold (*Dictyostelium discoideum*) | Wan et al., 2015 | **PRIDE:** PXD002320 |
| Fly (*Drosophila melanogaster*) | Wan et al., 2015 | **PRIDE:** PXD002321 |
| Mouse (*Mus musculus*) | Wan et al., 2015 | **PRIDE:** PXD002323 |
| Sea anemone (*Nematostella vectensis*) | Wan et al., 2015 | **PRIDE:** PXD002324 |
| Sea urchin (*Strongylocentrotus purpuratus*) | Wan et al., 2015 | **PRIDE:** PXD002325 |
| African clawed frog (*Xenopus laevis*) | Drew et al., 2020; Wan et al., 2015 | **PRIDE:** PXD017650, PXD017659; PXD002326 |
| Yeast (*Saccharomyces cerevisiae*) | Wan et al., 2015 | **PRIDE:** PXD002327 |
| **SOFTWARE AND ALGORITHMS** |  |  |
| Orthogroup inference | eggNOG v2.0.5 | https://github.com/eggnogdb/eggno-mapper/releases |
| Reference database construction | This paper | https://github.com/marcottelab/leca-proteomics/scripts/concat_ortho_proteins.py |
| Peptide identification | MSblender | https://github.com/marcottelab/MSblender |
| Ancestral genome inference | Count | http://www.iro.umontreal.ca/~csuros/gene_content/count.html |
| Feature extraction (CFMS data) | McWhite et al., 2020 | https://github.com/marcottelab/protein_complex_maps/tree/master/protein_complex_maps/features/ExtractFeatures/canned_scripts/extract_features.py |
| Feature integration (APMS data) | hu.MAP 2.0 | https://humap2.proteincomplexes.org/download |
|  | This paper | https://github.com/marcottelab/leca-proteomics/notebooks/map_entrez_to_eggno.ipynb |
| Pairwise protein interaction labels | This paper | https://github.com/marcottelab/leca-proteomics/scripts/label_featmat.py |
| Model optimization | TPOT 0.11.7 | http://epistasislab.github.io/tpot/ |
|  | This paper | https://github.com/marcottelab/leca-proteomics/scripts/run_tpot.py |
| Feature/model selection | This paper | https://github.com/marcottelab/leca-proteomics/scripts/select_features.py |
| Final model generation/assessment | This paper | https://github.com/marcottelab/leca-proteomics/scripts/predict_ppis.py |
| Protein complex detection | This paper | https://github.com/marcottelab/leca-proteomics/scripts/detect_communities.py |

### LECA gene set and resources

#### Resources for inferring the LECA gene set

Reference proteomes for 156 species (122 eukaryotes, 7 archaea, 27 bacteria; see Zenodo repository) were downloaded from the UniProt database along with the corresponding reference species tree^138^ for the parsimony analysis (**Figure S5**). This species tree and set of organisms were selected because they span the tree of life and serve as the gold standards curated by the Quest for Orthologs group for benchmarking orthology inference^139^. The species tree was downloaded from SwissTree (https://swisstree.sib.swiss/cgi-bin/swisst). Analysis of UniProt database reviewed proteins annotated with subcellular localizations was performed using the standardized SL accessions (see Zenodo repository), extracted with REST API queries.

#### Orthology mapping

Protein sequences from each reference FASTA file were searched against the eggNOG 5.0 database^38^ and mapped to orthologous groups (OGs) at the rootNOG level (taxonomic level = 1) using eggNOG-mapper v2.0.5^140^ with DIAMOND and a hit cut-off e-value of 10^-3^. As a result, 89,955 unique OGs spanning 156 species across the tree of life were used as input to the Dollo parsimony analysis. The group of rootNOGs assigned to the LECA node as a result of the parsimony procedure were converted to euNOGs (taxonomic level = 2759) with a set of hierarchical mapping files provided by Dr. Jaime Huerta-Cepas, the author of the eggNOG algorithm, *via* personal correspondence.

#### Dollo and Wagner parsimony

As no single model is assumed to be correct, instead, we use agreement among complementary reconstruction approaches to define a set of ancestral gene families for downstream biological interpretation. First, using the Count evolutionary analysis software^141^, we implemented a Dollo parsimony approach^39^ across 156 organisms to obtain an approximation of the LECA proteome. The Dollo parsimony model relies on the simplifying assumption that gene loss is irreversible, e.g., once a gene is lost it cannot be regained in a lineage. Thus, we determined the ancestral LECA proteome as the set of orthogroups either (a) shared by the respective outgroups (prokaryotes) and at least one of the eukaryotic species or (b) shared by two eukaryotic groups whose last common ancestor was LECA as defined by the gold standard species tree^142^. This approach has previously been shown to be effective at reconstructing likely LECA orthogroups^19^. However, there are caveats to its use, as detailed in the section below titled “Challenges and limitations to defining the LECA gene set”. Given these caveats, we additionally used the Count package to implement Wagner parsimony^143^ on the same organism/gene set as for Dollo. The Wagner model removes the irreversibility of Dollo gene loss events and allows for both gains and losses, thus accommodating horizontal gene transfer events.

#### Phylogenetic reconciliation

To complement the Dollo and Wagner parsimony–based inference of ancestral orthogroups, we implemented a large-scale phylogenetic reconciliation pipeline using Notung 3.0^144^ to more directly infer gene duplications, losses, and horizontal transfers relative to the reference species tree. We retrieved 10,089 eukaryotic-level gene trees (taxonomic level = 2759) from the eggNOG 5.0 database. For consistency with Notung input requirements, we generated a species-level taxonomic mapping by collapsing strain- and subspecies-level identifiers to their corresponding parent species and renaming each gene-tree leaf with its species mnemonic.

Following preprocessing, each gene tree was subjected to a stepwise Notung workflow consisting of (i) pruning to remove species absent from the reference tree, (ii) duplication–loss (DL) rooting, (iii) DL rearrangement of branches at 70% and 90% support thresholds, (iv) DL-rooting of the rearranged trees, and (v) duplication–transfer–loss (DTL) reconciliation with the reference tree under the “--phylogenomics” mode to infer orthogroup presence or absence across ancestral nodes. All procedures were executed in automated Bash and Python scripts (available on the supporting Github repository); final DTL-reconciled trees and tabulated origin/fam-present summaries are available on the supporting Zenodo repository. The resulting reconciliations yielded two parallel datasets (DTL70, DTL90) that were then summarized to identify orthogroups inferred to originate in LECA and to compare reconciliation-based with Dollo- and Wagner-based ancestral reconstructions. 1,295 (13%) of the OGs were not tested by reconciliation, largely due to species differences between the eggNOG gene trees and the reference species tree.

#### Investigation of orthologous groups of unknown function

We attempted to assign functions, or, at the minimum, subcellular localizations, to the 25% of uncharacterized (by eggNOG) LECA OGs using the UniProt database. To this end, we downloaded 363,430 proteins from the UniProt database that were (a) “reviewed” status and (b) assigned a standardized subcellular localization (SL) ID by Uniprot for each compartment that we trace back to the last eukaryotic common ancestor (**Figure S2A**). The total number of proteins assigned to a subcellular compartment varied by four orders of magnitude, where 166,296 proteins were assigned to the cytoplasm at the highest end and 116 proteins were assigned to phagocytic cups at the lowest end (**Figure S2A**). Furthermore, we investigated the diversity and magnitude of eukaryotic and prokaryotic species contributing to these annotations (**Figure S2B**) and observed an underrepresentation of clades outside Amorphea (see tree in **Figure S5** for supergroup organization). For example, the nucleus is considered a distinguishing feature of eukaryotes; UniProt proteins annotated to localize to the nucleus come predominantly from 1,032 distinct Amorphean species, with almost two orders of magnitude fewer such annotations contributed by non-amorphean species, consisting of 152 archaeplastidans, 2 cryptophytes, 21 excavates, and 62 TSAR species.

To quantify what proportion of the LECA gene set is represented in UniProt, we mapped all 363,420 reviewed UniProt proteins mentioned above to eukaryotic orthologous groups (“euNOGs”; NCBI taxonomic identifier = 2759). This resulted in 13,556 unique euNOGs, the percentage of which that trace back to LECA varies significantly by subcellular localization (**Figure S2C**). As an aside, we note that the same euNOG can often be assigned multiple UniProt SL IDs, netting a total 24,628 euNOG-SL mappings.

Of the 2,790 ciliary proteins that map to 478 unique eukaryotic orthologous groups (euNOGs), nearly ∼25% of the 299 UniProt ciliary euNOGs that intersect with LECA OGs were originally assigned “unknown function” by the eggNOG functional annotation algorithm–a larger proportion than most of the other eukaryotic compartments described within the UniProt database. Similarly, of 1,317 cytoskeletal euNOGs, ∼60% trace back to LECA and 111 of those were assigned the “function unknown” eggNOG category. Thus, the combination of eggNOG and Uniprot annotations provided a reasonable initial annotation set for subsequent analyses.

#### Challenges and limitations to defining the LECA gene set

Binning proteins into evolutionarily related orthologous groups with respect to the root of the eukaryotic tree nets a “coarse-grained” mapping of the relationships between eukaryotic genes, *i.e*., we can not rigorously distinguish orthologs from paralogs, which could limit the degree to which we can infer functional continuity across homologous gene, as orthologs are argued to be more likely to retain ancestral functions than paralogs (and xenologs), although such trends have generally been weak when measured^145,146^. With that said, we are still able to draw conclusions about the properties of families of genes rather than the pairwise relationships of individual members; this approach is intuitive and convenient for large-scale systematic studies and broadly supported^147–149^, albeit not resolving the phylogenetic relationships among the genes within each orthogroup. However, there is considerable disparity between OG assignment algorithms, though eggNOG has been demonstrated to have among the highest accuracies when tested on a benchmark set of manually curated orthologs^19^. In the same study, the eggNOG algorithm was also shown to perform best at detecting distant homology and properly splitting out-paralogs, making it the best suited algorithm currently available for our goals. Some protein families and eukaryotic lineages with fast rates of evolution (e.g., transcription factors, proteins associated with the innate immune response, and in general plants that are prone to whole genome duplication^150–154)^ remain a weakness to the approach. Proteins such as these are likely “under-split” with respect to their associated orthologous groups. Leucine-rich repeat proteins are a salient example: more than 100 human LRR proteins were assigned to KOG0619, an OG we traced back to LECA related to intracellular trafficking and secretion, and this trend persists across nearly all eukaryotes sampled that had proteins assigned to KOG0619. In this way, we are most likely underestimating the size of the distinct LECA gene set.

**Table S1** reports the observed support under each method for every LECA OG, so that researchers can check the level of support for specific gene families of interest. Here, we note the strengths and weaknesses of each method contributing to the LECA OG determination. First, the Dollo parsimony procedure used to bound the overall LECA OG set assumes that the probability that a trait emerges more than once is negligible ^39^ and is the simplest form of ancestral state inference. The use of Dollo parsimony is justified as a base model^155–157^ given that (a) our goal was to determine a binary character state (the presence or absence of genes), (b) we had a consensus reference species tree in hand^138^, (c) the target gene set is eukaryotic wherein independent gene losses are common and gains of multiple genes are (relatively) rare, (d) the expected influence of horizontal gene transfer is thought overall to be minimal (estimated to be ∼1% of genes or less^158^, but see below), and (e) probabilistic ancestral state reconstruction methods, such as phylogenetic birth-death-gain models, are prohibitively slow for a data set of this size. Nonetheless, one flaw in this approach is particularly worth noting: multiple species within Excavata host plastids or plastid-derived genes orthologous to plastid proteins in plants^159–161^, even though it is widely accepted that primary plastids share a single origin^162,163^ and Archaeplastida is monophyletic^164,165^. If the Archaeplastidan monophyly is to be believed^166,167^, the last common ancestor of Excavata independently acquired plastids (violating Dollo’s law), resulting in the inflation of our LECA gene set by ∼40 plastid-associated OGs. To correct for this error, we manually removed these OGs from consideration during construction of the LECA interactome.

Due to these limitations with Dollo, we applied additional phylogenetic procedures to each of the Dollo-estimated LECA OGs in order to annotate any additional support by these algorithms. Importantly, as the true extent of horizontal gene transfer in early eukaryotes is not well determined, we used methods that considered the possibility of horizontal gene transfer events, Wagner parsimony and duplication-transfer-loss (DTL) phylogenetic reconciliation. Of the 10,091 LECA OGs from Dollo, 6,429 were additionally supported by either Wagner or DTL reconciliation. Notably, an additional 582 OGs were reconciled as Eukaryotes not Excavata. While some of these cases may simply reflect limited taxon sampling in the selected species tree, it is worth noting that a recent tree places the root of the eukaryotic tree within Excavata^168^, and the “Eukaryote not Excavata” OGs may still represent potential LECA OGs. For OGs conserved across only a subset of major clades, changes to the species tree used for reconciliation might thus impact estimates of OG ages, although the most broadly conserved OGs should be reasonably robust to this effect. For the final set of 3,193 OGs present in the estimated LECA interactome, 98% are supported by either Wagner or DTL reconciliation, indicating robust support for their presence in LECA, at least under the caveats discussed here.

Finally, errors in gene trees (as well as reliance on individual estimated gene trees) can degrade performance in reconciliation methods; these methods have also been observed to exhibit higher false positive rates for inferred horizontal gene transfer events as compared to implicit phylogenetic methods such as Count’s Wagner parsimony implementation (e.g. ^169^). Hence, these methods can also be subject to age estimation errors, potentially favoring younger ages.

### Resources for interactome mapping

Biological samples, mass spectrometry data sets, and software used in this analysis are summarized in the **Resources Table**. Proteomes for 31 eukaryotic species were sourced as summarized in **Table S3**.

### Mass spectrometry

#### Native protein extraction and fractionation

For lysates described below protease inhibitor cocktail was cOmplete mini EDTA-free (Roche), phosphatase inhibitors were PhosSTOP EASY pack (Roche), and all steps after addition of lysis buffers were conducted at 4°C or on ice unless otherwise indicated. Native soluble extracts were quantified by DC Protein Assay (BioRad). All protein samples were 0.45 µm filtered (Ultrafree-MC-HV Durapore PVDF, Millipore) prior to chromatography. Chromatography was performed on an HPLC system as in ^33^ unless otherwise stated.

*Brachionus rotundiformis* was collected in batches on Filter Mesh 100 Nylon (∼65 µm pore) to remove feeder algae prior to flash freezing in liquid nitrogen. Frozen material (3.1 g) was ground to power in a liquid nitrogen-chilled mortar and pestle and resuspended in an equal volume of *Tetrahymena* Lysis Buffer (25 mM Tris pH7.4, 25 mM NaCl, 1 mM EDTA, 10 % glycerol, 0.2% NP40, with 1 mM DTT, 1 mM PMSF, phosphatase inhibitors, and protease inhibitor cocktail added freshly). Cells were disrupted with 10 strokes in a glass dounce fit with a tight pestle. Following centrifugation 3000 x g, 10 minutes to remove debris, the supernatant was clarified twice by centrifugation 20,000 x g 10 minutes. Size Exclusion Chromatography was performed with 2.6 mg extract in a 200 µl sample loop and mobile phase Buffer S (50 mM Tris-HCl pH 7.5, 50 mM NaCl).

*Phaeodactylum tricornutum* (UTEX 646) grown without silica was briefly washed by pelleting (2000 x g, 10 minutes, 21°C, no brake) and resuspended in 0.5x artificial seawater (UTEX) before collecting (3000 x g, 4°C, slow deceleration) and flash freezing. Frozen material was ground to powder and allowed to thaw before refreezing and regrinding. 1g of powdered material was resuspended in 800 µl Lysis Buffer (50 mM Tris pH 7.5, 150 mM NaCl 5 mM EGTA, 10% glycerol, 1% NP40 with 0.1mM DTT) with phosphatase inhibitors and Plant Specific Protease Inhibitors (Sigma # P9599). Material was frozen and thawed again before sonicating 6 x 10 seconds on, 20 seconds off, 70% duty cycle. Lysis was monitored by microscopy. The extract was incubated on ice with periodic gentle vortexing for 30 minutes prior to clarification twice at 14,000 x g, 10 minutes. Extract was diluted 3-fold with 50 mM NaCl prior to loading 2 mg for SEC separation as above. For separation by mixed bed ion exchange chromatography (Poly CATWAX A, PolyLC Inc.) salt was reduced by 5x dilution with 10 mM Tris pH 7.5, 5% glycerol, 0.01% NaN_3_ and proteins were re-concentrated by ultrafiltration (Amicon Ultra 0.5 ml 10,000 MWCO). IEX chromatography was with 1.9 mg in a 250 µl sample loop.

*Euglena gracilis* (UTEX 753) was washed briefly by centrifugation (1,500 x g, 5 minutes, 21°C) and resuspension in dH_2_0, before collection by centrifugation and flash freezing. Material was ground as above and 4.7 g was resuspended in Lysis buffer plus both the cOmplete mini EDTA-free protease inhibitors and the Plant-Specific Protease Inhibitors (Roche). Lysate was sonicated 9 x 10 seconds on, 20 seconds off, 60% duty cycle, followed by gentle nutation 30 minutes. Debris was removed by centrifugation 1,500 x g, 10 minutes, and the supernatant was further clarified twice with 14,000 x g, 10 minute spins. Final extract was filtered through a 0.45 µm syringe filter (Durapore PVDF, Millipore) prewashed with dH_2_0. Extract was diluted 4-fold in Buffer S and 2 mg loaded on a 200 µl sample loop for SEC fractionation.

Two fresh pig tracheas (*Sus scrofa*) were shipped on ice from Sierra for Medical Science arriving within 24 hours of harvest. After removal of fat tissues the trachea were slit lengthwise, chopped crosswise into several pieces, and washed with multiple changes of ice cold PBS pH 7.4 to remove serum and blood cells prior to extraction with 100 ml Ca++ shock buffer as in ^170^ including protease and phosphatase inhibitors at 0.5x concentration and 0.1 mM PMSF. Cilia were released by vortexing and manual agitation for 10 minutes. Debris was pelleted 500 x g, 2 minutes and floating lipids were removed by aspiration. Cilia were collected by centrifugation 12,000 x g 10 minutes and washed once by resuspension and centrifugation. Ciliary pellets were resuspended in Ca++ shock buffer with 1% NP40 to extract soluble proteins and residual axonemes were removed by centrifugation twice at 12,000 x g 10 minutes. Any floating lipids were removed after each spin. Extract was flash frozen until used. Thawed extract was diluted 2-fold with 10 mM Tris pH 7.5, 5% glycerol, 0.01% NaN_3_ and re-clarified 12,000 x g 10 minutes prior to ultrafiltration with 30,000 MWCO Ultracel Amicon Ultra 0.5 ml units to load 1.7 mg in a 250 µl sample loop for IEX chromatography.

*Tetrahymena thermophila* SB715 were grown and cilia extracts made as in ^171^ except that deciliation was by pH shock according to ^172^. 1.5 mg cilia extract was fractionated by mixed bed IEX and 1.2 mg by SEC with SEC mobile phase Buffer S-C (50 mM Tris-HCl pH 7.4, 50 mM NaCl, 3 mM MgSO_4_, 0.1 mM EGTA). Deciliated *Tetrahymena* “bodies” were collected by centrifugation 1,700 x g, 5 minutes, washed once by resuspension in Deciliation Medium (10 mM Tris-HCl pH 7.4, 10 mM CaCl_2_, 50 mM sucrose), collected by centrifugation as before and flash frozen until use. *Tetrahymena* body lysate was prepared by liquid nitrogen grinding frozen material before resuspending in an equal volume *Tetrahymena* Lysis Buffer with 0.1 mM PMSF. Lysis was achieved on ice for 10 minutes by pipetting up and down. Debris was removed by centrifugation 3,000 x g, 10 minutes. Supernatant was clarified and floating lipids removed by sequential centrifugations at 40,000 x g, 10 minutes, 45,000 x g 30 minutes, 130,000 x g 1 hour, and 130,000 x g 45 minutes. Extract was diluted (final NaCl 22 mM) prior to loading 2.2 mg on a 250 µl sample loop for IEX chromatography. The remaining extract was flash frozen and thawed later for SEC chromatography. Extract was clarified 25,000 x g 10 minutes immediately after thawing, and again after dilution in Buffer S-C for loading of 1.4 mg in 200 µl sample loop. For DSSO-crosslinked samples, cilia extract was prepared using the pH shock method as above, but 20 mM HEPES pH7.4 was substituted for the 50 mM Tris of the Cilia Wash Buffer. Extract was concentrated by ultrafiltration in an Amicon Ultra Ultracel 10k NMWL unit (UFC501096) to load 1.5 mg in a 250 µl sample loop. The final concentration of NP40 was 2.75%. Fractionation on a mixed bed IEX was performed with substitution of 10 mM HEPES pH 7.4 for Tris in the chromatography buffers A and B. For crosslinking DSSO was dissolved freshly in dry DMF to 50 mM and then diluted with 10 mM HEPES pH 7.4 to 10.5 mM before dispensing 25 µl into each 500 µl fraction. To ensure activity of the crosslinker the DSSO solution was prepared in 2 consecutive batches to treat a total of 76 column fractions. Crosslinking proceeded 1 hour at room temperature (∼21°C) and was quenched by addition of Tris pH 8.0 to 28 mM.

*Xenopus laevis* sperm were isolated from dissected testes of five or eight J-strain *Xenopus laevis* males. Testes were perforated with a 25-gauge needle, sperm blown out using MMR (Marc’s Modified Ringers). Larger debris was allowed to settle, and liquid transferred to a fresh tube. Sperm were collected by centrifugation 1,500 x g, 10 minutes. Supernatant was discarded and the sperm pellet was lysed by resuspension in an equal volume of Sperm Lysis Buffer (10 mM Tris-HCL pH7.5, 20 mM KCl, 5 mM MgCl_2_, 5% glycerol, 1% n-Dodecy-ß-D-Maltoside (Anatrace) with 0.5 mM DTT added freshly). Lysate was clarified by centrifugation 14,000 x g, 10 minutes. 1.2 mg was loaded for mixed bed IEX column fractionation (PolyLC Mixed-Bed WAX-WCX, PolyLC Inc. #204CTWX0510), and 3 mg for SEC fractionation (BioSep-SEC-s4000, Phenomenex).

*Mus musculus* (embryonic stem cells) were grown as described in ^173^. Cells were harvested without trypsin by washing in ice cold phosphate buffered saline (PBS), pelleted, and placed on ice. A 250 µl cell pellet was lysed on ice (5 min) by resuspension in 500 µl of Pierce IP Lysis Buffer (25 mM Tris-HCl pH 7.4, 150 mM NaCl, 1 mM EDTA, 1% NP-40 and 5% glycerol; Thermo Fisher) containing 1x protease inhibitor cocktail III (Calbiochem). During the 5 minutes, cells were periodically dounce homogenized with a small-clearance glass pestle (pestle B). Approximately 2 mg of total protein was loaded on either a mixed bed IEX column (PolyLC Mixed-Bed WAX-WCX, PolyLC Inc. #204CTWX0510) or a BioSep-SEC-s4000 gel filtration column (Phenomenex) equilibrated in PBS, pH 7.2. HPLC chromatography was as in ^33^ and collected fractions were processed as described in ^173^.

Lysate preparation and chromatographic separation of other species is described in ^29,33,171,174^.

### Data acquisition and processing

All column fractions were reduced, alkylated and digested with trypsin for mass spectrometry by either method 1 or 2 of the protocols in ^175^. Spectra were collected as in^33^ on either a Thermo Scientific Orbitrap Fusion Tribrid or an Orbitrap Fusion Lumos Tribrid mass spectrometer except as noted below. Euglena data were collected on a Lumos using CID (35%) and a topspeed 75 minute method as in ^33^. Spectra for DSSO crosslinked *Tetrahymena* cilia IEX fractions were collected using a 2 hour DDA MS2-MS3 method as described in ^171^ but processed for protein identifications in this study using the MSBlender pipeline described below.

### Computational analyses

#### Reference database construction

Protein sequences from each of the 31 reference FASTA files in **Table S3** were compared against the eggNOG 5.0 database^38^ and mapped to orthologous groups (OGs) at the euNOG level (taxonomic level = 2759) using eggNOG-mapper v2.0.5^140^ with DIAMOND and a hit cut-off e-value of 10^-3^. Using a previously well-tested strategy for comparative proteomics^33^, for each species, a reference database was constructed where proteins are binned into their respective OGs such that each FASTA entry represents a bin of proteins or protein family; this was accomplished by concatenating each sequence assigned to an OG interposed with a triple lysine sequence. Since we allow for two missed trypsin cleavages in peptide spectra assignment, this triple lysine sequence ensures that we avoid the misassignment of peptides matching a chimera of two binned sequences. The benefits of this approach are three-fold: (1) defining proteomes in terms of OGs enables cross-species comparisons, (2) OG binning recovers peptide mass spectra that otherwise could not be uniquely assigned to highly sequence-similar proteins, and as a natural extension (3) facilitates proteomic analysis of species with high ploidy, e.g., *X. laevis* (allotetraploid)^174^ and *T. aestivum* (allohexaploid)^33^.

#### Peptide mass spectra processing

Matching of mass spectra to peptides was performed with MSGF+, X!Tandem, and Comet-2013.02.0, each run with 10ppm precursor tolerance and allowing for fixed cysteine carbamidomethylation (+57.021464) and optional methionine oxidation (+15.9949). Peptide search results were integrated with MSBlender ^176^ as described in ^29,33^ with the exception that high confidence (1% FDR) peptide spectral matches were required from two out of the three peptide identification algorithms. In all, we measured 379,758,411 peptides that were uniquely assigned to 259,732 unique proteins and orthogroups across all fractions. These results were filtered such that we only retain orthogroups that (a) were determined to trace back to the last eukaryotic common ancestor and (b) were strongly observed such that the sum total peptide spectral matches (PSMs) across all fractionations was >= 150.

#### Feature curation for protein-protein interactions

For each orthogroup found in each MS fractionation for each species sample, an elution vector was constructed by concatenating the peptide spectral counts for each orthogroup in each fraction. Four measures were used to compare all pairwise elution vectors: the Pearson correlation coefficient, Spearman’s correlation coefficient, Euclidean distance, Bray-Curtis dissimilarity. These measures were computed as described in ^175^ and were generated for: (1) vectors for 149 individual fractionations, (2) concatenated vectors that include all samples within the Amorphea eukaryotic supergroup, (3) concatenated vectors that include all samples within the Excavate eukaryotic supergroup, (4) concatenated vectors that include all samples within the TSAR eukaryotic supergroup, (5) concatenated vectors that include all samples within the Archaeplastida eukaryotic supergroup, and (6) concatenated vectors that include all eukaryotic samples, netting 616 CFMS features.

In order to specifically target conserved pan-eukaryotic protein interactions, we required elution vectors for each protein-protein interaction (PPI) to have a minimum Pearson *r* of 0.3 and be observed in at least two of the four eukaryotic supergroups, *i.e.*, Amorphea, Excavata, TSAR, and/or Archaeplastida (**Figure 2B**). This reduced the size of our input data from 17,895,154 pairwise protein comparisons to a curated set of 4,491,719 highly conserved PPIs. Finally, we integrated the intersection of these conserved PPIs with 47 pairwise features generated from an orthogonal collection of ∼15,000 mass spectrometry proteomic experiments^30^ that include APMS^53–55^, proximity labeling^56,57^, and RNA-pulldown data^58^ to attain our final PPI feature matrix, resulting in a total of 663 features for each of 4,491,719 highly conserved potential pairwise PPIs.

#### Assembly of gold standard protein complexes

Gold standard protein interactions were downloaded from the CORUM^59^ (http://mips.helmholtz-muenchen.de/corum) and Complex Portal^60,61^ (https://www.ebi.ac.uk/complexportal) databases. Both databases include protein-protein interactions for multiple species, spanning multiple mammals in CORUM (human, rat, mouse, cow, pig) and many eukaryotes in Complex Portal (human, rat, mouse, cow, pig, yeast, *Arabidopsis*, worm, fly, chicken, snake, fish, frog, rabbit). Redundant complexes were merged, and (to reduce representational bias) any complex with >30 subunits was removed from the gold standard complex set. Finally, UniProt IDs were matched to euNOG IDs and the gold standard complexes were pruned to only include those in the LECA proteome as determined by the ancestral state reconstruction described above.

#### Machine learning for protein interactions

All gold standard PPIs observed in our filtered data set were labeled as positive interactions. Negative interactions were defined as interactions between proteins in different gold standard complexes (e.g., given two gold standard heterotrimers A-B-C and X-Y-Z, data corresponding to an A-X protein pair would be labeled as a negative interaction). To mitigate class imbalance, the total number of negative labels was limited to 3X the observed number of positive PPIs in our data, resulting in 6,629 total positive PPI and 19,887 total negative PPI labels in our feature matrix.

Positive PPIs that participate in multiple complexes are a potential source of representation bias in the truth set, which can lead to under or overfitting during model training. To overcome this, we implemented a data stratification approach (**Figure S6**). First, all gold standard protein complexes are given a unique numeric ID. All protein pairs within a complex inherit that ID. Negative PPIs receive group labels by randomly sampling the distribution of positive PPI group IDs with replacement. If a protein pair participates in >1 complex, that pair will be labeled with a list of IDs. These ID lists represent networks of overlapping complexes. We implemented transitive closure of the networks by recursively merging ID lists that overlap with each other, netting a fully stratified “supergroup” label for each gold standard PPI in the data set.

A group-based split method (scikit-learn’s “GroupShuffleSplit” class) was used to generate 5 sets of test and training data, where 75% of the labeled data was used for training and 25% for testing. Each of the 5 training sets were used as input into TPOT^177^, an automated machine learning pipeline built on top of scikit-learn, to find the best classification method, pre-processing steps, and parameters for our data. While TPOT generates an internal cross-validation score to evaluate the performance of different models, the pipeline is agnostic to strata within the training set and is thus subject to overfitting. In each instance, the 25% hold-out set was used as a true test of the optimized models produced by TPOT. These results are reported in **Table S4**. In the majority of cases, TPOT reports the extremely randomized trees (ExtraTrees) algorithm with slightly varying parameters and pre-processing steps as the best model for our data. However, both a stochastic gradient descent and linear support vector classification (SGDClassifier and LinearSVC, respectively) pipeline scored comparably to the ExtraTrees method, so we moved forward with feature selection and model assessment for those pipelines as well.

To further reduce the risk of overfitting, recursive feature elimination with cross-validation (RFECV) was used to obtain feature importances and an optimal feature set for each of the chosen models (see Zenodo data repository). Feature importance was evaluated using either the Gini index (in the case of ExtraTrees) or the absolute value of the coefficients of the linear model (in the case of LinearSVC and SGDClassifier). While RFECV generates an internal cross-validated test score to determine an optimal feature set, the module is agnostic to stratified data and is thus subject to sampling bias. Again, we employed a custom group-based split approach (scikit-learn’s “GroupKFold” class, illustrated in **Figure S7**) to generate 5 sets of test and training data. Since GroupKFold generates test/train splits such that every protein complex supergroup is included in the test data at least once, this allows us to use hold-out test sets to gauge bias in the “best” feature sets output by RFECV while also evaluating feature importance stability.

RFECV determines an optimal feature set following these steps: (1) the input training data is used to fit a given model, resulting in either a Gini index value or coefficient for each feature in the data set; (2) the least important feature(s) are recursively removed and the mean test accuracy is computed using cross-validation across the input training set, (3) the optimal number of features is determined such that mean test accuracy is maximized. Then, we use the holdout test set to evaluate the true performance of the final model as output by RFECV method. True feature importance was assessed by aggregating the RFECV results of each split, and, for each feature, counting the number of times it appears in the “optimal” set output and computing the mean and relative standard deviation of its Gini index/coefficient. The “best” features are those that maximize the number of appearances in the final “optimal” set across GroupKFolds splits, maximize the absolute value of the Gini index/coefficient and minimize relative standard deviation (*i.e*., low RSD indicates the feature importance is stable across GroupKFold train/test splits).

We measured precision and recall for each model using the top 5, 10, 25, 50, 100, and 250 highest ranked features (determined on a model-by-model basis, in other words, the “top” features for the LinearSVC classifier are different than the “top” features for the ExtraTreesClassifier) and all 663 features (**Figure S8A**), choosing the feature set that had the highest number of PPIs and unique proteins within a 10% FDR threshold (**Figure S8B,C**).

#### Community detection of protein complexes

Communities of interacting proteins were identified using the walktrap algorithm *via* the igraph library interface in Python. Briefly, the walktrap algorithm detects community structure by executing a user-defined number of random walks from each vertex in a graph. Random walks tend to become “trapped” in strongly connected sub-communities of the graph^62^, a behavior that we reinforce by weighting graph edges with the probability scores output by our three PPI models (LinearSVC, ExtraTrees, and SGDClassifier). Then, individual protein complex clusters are partitioned such that the modularity of the network is maximized^178^, netting an “optimal” number of protein complex communities given the input graph structure. The walktrap algorithm performed best with the PPI scored output by the LinearSVC model, and was thus chosen as our final interactome. The features used in the final model are reported in the Zenodo repository.

#### Validation of the protein interaction model with external data

External interaction datasets were sourced from ^63–65^. The likelihood of LECA PPIs (*I_L_*) agreeing with external interaction networks (*I_E_*) was calculated with a formula analogous to an odds ratio, described by the equations below.

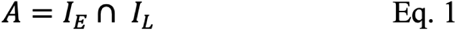

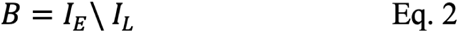

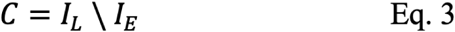

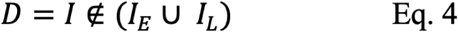

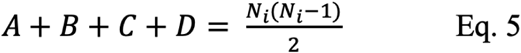

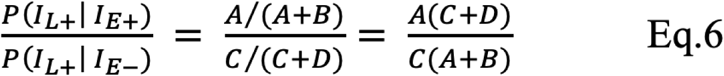

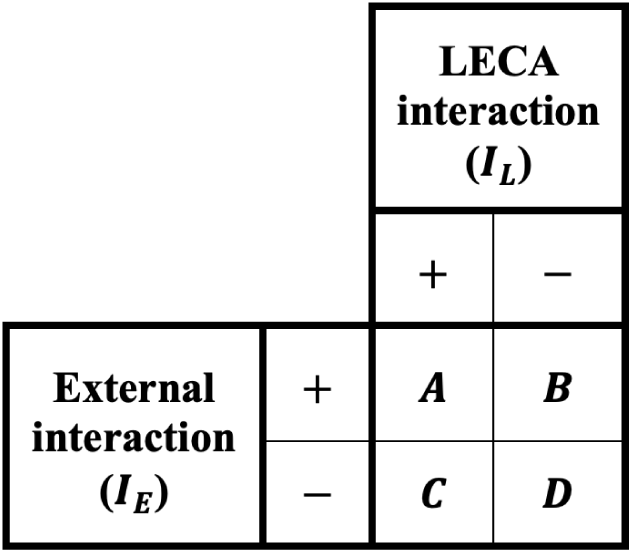

### Challenges and limitations to mapping the LECA protein interaction set

It is important to note that the data used in this study strongly favors humans and mammals in general. Most of the CFMS and APMS experiments in this study are sourced from humans. Of the 25,000 experiments included in this study, approximately 18,000 are derived from human cells. Gold standard protein complexes obtained from the CORUM database are exclusively mammalian, which motivated us to also incorporate gold standard PPIs from the ComplexPortal database. However, though ComplexPortal is more diverse and includes *Arabidopsis* assemblies, the majority of the data is still Amorphean. We took a number of steps to ensure we target pan-eukaryotic proteins and protein interactions; for example, we filtered the MS data to only include proteins that we trace back to LECA as well as require every PPI be observed and reasonably correlated by CFMS (Pearson *r* > 0.3) in at least 2 of the 4 eukaryotic supergroups (as defined in **Figure 2B**) prior to entry into the machine learning pipeline, with this latter requirement helping to ensure that PPI support is not drawn solely from opisthokonts.

Ideally, it would be preferable to experimentally determine interactions independently in each species and use these as traits for evolutionary analysis. In practice, however, such an approach is deeply underpowered, as all current protein interaction mapping technologies, including CF-MS, affinity-purification / MS, immunoprecipitation / MS, yeast 2-hybrid, and even computational approaches such as AlphaFold, exhibit high false negative rates for detecting and measuring protein interactions^179^. This tendency is exacerbated further by the limited sampling of tissues and cell types imposed by a study of this scale, hence a full evolutionary analysis is not sufficiently powered to be effective with current datasets. This limitation motivates the current (better powered) approach to determining interactions as accurately as possible using the pooled data across species, while requiring supporting evidence from multiple supergroups to ensure a high probability of the interactions being ancestral.

Co-fractionation mass spectrometry identifies stable complexes that survive biochemical fractionation. Stochastic sampling across a large number of species and fractionations allows us to recover some transient interactions. CFMS is also biased towards abundant and soluble proteins, though we typically employ detergents to improve coverage of membrane proteins. We measure approximately 60% of the 10,091 LECA OGs (probably due to the above detection biases) and high precision PPIs for half of these, indicating a high false negative rate. Inclusion of the APMS data sets increases the power of our model for PPI detection but only for systems conserved within Amorphea. Binning proteins into evolutionarily related protein families (orthogroups) prior to peptide identification and assignment results in loss of resolution for different isoforms and some paralogs. For the most part, we are able to draw conclusions about the properties of families of genes rather than individual members. In some cases, we can retroactively disentangle which variant(s) or paralog(s) participate in a specific interaction by examining the specific peptides identified by mass spectrometry. Moreover, the loss of family member resolution is offset by improved protein identifications: Since the proteomes must be defined in terms of OGs in any event in order to effectively perform cross-species comparisons, by moving that process to the protein identification step, we additionally benefit from the increased protein identification rates by allowing orthogroup-restricted peptides to contribute to identification and quantification, significantly improving proteome coverage (see ^33^ for quantification of this effect).

### Resources for disease analyses

We downloaded 17,019 gene-disease relationships from the Online Mendelian Inheritance in Man database (omim.org)^119^.

### Curation of a gene-disease data set

Ensembl accessions from the OMIM data set were first mapped to human UniProt identifiers and then subsequently matched with their corresponding eggNOG orthologous groups (OGs) at the root of eukaryotes (NCBI taxonomy level = 2759). Gene-disease assignments in the raw OMIM data did not follow a standardized schema and required a combination of programmatic and manual cleaning. For example, AKT1, PTEN, KLLN, and SEC23B are respectively assigned to Cowden syndrome 6, Cowden syndrome 1, Cowden syndrome 4, and Cowden syndrome 7 in the original data set. After cleaning, these genes are grouped under a common “Cowden syndrome” label. We filtered the data set to contain only LECA OGs, netting 5,761 genotype-to-phenotype associations for the network propagation of gene-disease relationships in the conserved eukaryotic interactome. In total, we curated 1,683 unique disease labels for 2,262 highly conserved human genes.

### Network propagation

We used a cross-validated network propagation approach to systematically assign disease predictions to proteins in the interactome. If a protein in the network is known to be associated with a disease, then each connecting node receives the score of the connecting edge. This process is repeated for each unique disease label in our OMIM data set given that the disease has at least five mapped associations with genes that also map to an orthologous group in the last eukaryotic common ancestor. To assess the quality of this propagation for each disease, we iteratively leave out true positive nodes and query how well the propagation recapitulates known gene-disease relationships (*i.e*., leave-one-out cross validation). We calculate true and false positive rates to construct a receiver operating characteristic (ROC) curve as a function of propagated score. Then, we use the area under the ROC curve (AUROC) as a measure of performance. Additionally, for each disease, we repeat propagation from randomly selected nodes from the network to evaluate the statistical strength of the gene-disease network versus randomly assigned gene-disease relationships.

### Analysis of yeast gene-mutant phenotype data sets

We downloaded results from 14,484 phenotypic screens across 4,554 mutant yeast strains *via* the YeastPhenome project^180^. Normalized phenotypic values (NPVs) are used in the YeastPhenome data set to measure gene-phenotype associations. NPVs greater than 3 standard deviations (s.d.) above the mutant strain mean or less than -3 s.d. below the mean denote significant gene-phenotype associations. This data set was split by NPVs into associations with NPVs greater than or equal to 3 (“upper”) and less than or equal to -3 (“lower”). This resulted in 226,050 upper gene-phenotype associations and 458,562 lower gene-phenotype associations, netting 684,612 significant associations in yeast across 19,607 phenotypes. The genes for these associations were matched to their corresponding LECA OGs, and these gene sets for each yeast mutant phenotype analyzed *via* network propagation in the LECA interactome as described above for the OMIM gene sets.

### Analysis of *Chlamydomonas* gene-mutant phenotype data sets

We downloaded reported phenotypic measurements for 747 *Chlamydomonas* mutant strains^181^ and selected all 1,636 significant (FDR < 0.3) gene-phenotype associations, spanning 167 unique phenotypes. Assayed genes were matched to their corresponding LECA OGs, then the gene sets for each *Chlamydomonas* phenotype analyzed *via* network propagation in the LECA interactome as described above for the OMIM gene sets.

### Experimental analyses of candidate disease genes

#### Genetic knock outs in Mus musculus

We sourced *Atp6v1a* knockout data (with permission) from the Knockout Mouse Program (KOMP2) sited at the Baylor College of Medicine, which resides under the umbrella of the International Mouse Phenotyping Consortium (IMPC). The IMPC aims to systematically phenotype mice that are homozygous for a single-gene knockout or heterozygous when homozygotes are lethal or sub-viable^182^. Within the IMPC, KOMP2 production centers use the high-throughput and rigorously standardized IMPReSS pipeline to generate and phenotype single-gene knockout mice. Methods for generating single-gene null alleles are described in ^183^ and phenotype data collection procedures can be accessed in detail at https://www.mousephenotype.org/impress/index.

*Atp6v1a* mutant alleles were generated by KOMP using CRISPR/Cas9 to introduce a critical exon deletion in a murine C57BL/6N background. Phenotypes were measured from postnatal mice following the embryonic and early adult IMPC pipelines. In the case of *Atp6v1a*, homozygous knockouts resulted in complete penetrance of pre-weaning lethality. As a result, heterozygous knockouts were generated for 8 female mice and 8 male mice. At 14 weeks, bone mineral content and density was measured for each *Atp6v1a*^em1(IMPC)Bay/+^ mutant using dual-energy X-ray absorptiometry (DEXA). The IMPC uses the PhenStat R package to identify abnormal phenotypes from high-throughput pipelines ^184^; for the *Atp6v1a*^em1(IMPC)Bay/+^ mutants, a linear mixed model factoring in the effects of sex and body weight was implemented in PhenStat to assess significance of differential bone mineral content.

#### EFHC2 patient genetics

Individual A4237-22 was a male of Egyptian origin who was diagnosed with small kidneys, increased echogenicity, cortical and medullary cysts, and microcephaly. To identify a potential genetic cause for the individual’s phenotype, authors S.S. and F.H. performed whole exome sequencing (WES) analysis on individual A4237-22. Given the parents’ unaffected status regarding their renal phenotype, a recessive mode of inheritance was hypothesized.

Homozygosity mapping revealed only 4.4 Mb of homozygosity, confirming the non-consanguinity of the parents (**Figure S4A**). We detected a hemizygous X-linked missense variant in A4237-22 (c.398G>A; p.Arg133His) (**Figure 6D, S4B**). The variant has not been reported as homozygously or hemizygously in the gnomAD database in 166,211 control individuals. The p.Arg133His amino acid resides in the DM10 domain (**Figure 6E**).

The patient’s DNA was also screened for potentially deleterious variants in all genes known to cause kidney disease without results.

#### Protein localization and knockdown experiments in Xenopus laevis

*Xenopus* embryo manipulations were performed as in ^185–187^. Briefly, female adult *Xenopus* were ovulated by injection of hCG (human chorionic gonadotropin). In vitro fertilization was carried out by homogenizing a small fraction of a testis in 1X Marc’s Modified Ringer’s (MMR). Embryos were dejellied in 1/3X MMR with 2.5% cysteine (pH 7.8) at the two-cell stage. For microinjections, embryos were placed in a 2% Ficoll and 1/3X MMR solution, injected with mRNA using forceps and an Oxford universal micromanipulator, and washed with 1/3X MMR after 2 hours.

The full length sequences of *Xenopus EFHC2* and *GLG1* were downloaded from Xenbase^188^. The DNAs corresponding to the open reading frames (ORFs) of *EFHC2* and *GLG1* were amplified from *Xenopus* cDNA and were cloned into a pCS10R MCC vector containing an N-terminal GFP or a C-terminal FLAG tag driven by an MCC specific alpha tubulin promoter, respectively. The pCS10R MCC *GFP-EFHC2* R133H construct was generated by site-directed mutagenesis (NEB, #E0554S) from pCS10R MCC *GFP-EFHC2*. Capped mRNAs were synthesized using the mMESSAGE mMACHINE SP6 transcription kit (Invitrogen Ambion, #AM1340). A morpholino antisense oligonucleotide (MO) against *GLG1* was designed to block translation (GeneTools). The MO sequence is 5’-CCATCTTGGGAAGTGCTAGTCAAG-3’.

mRNA and MO were injected into two ventral blastomeres of 4-cell stage Xenopus embryos in 2% Ficoll (w/v) in 1/3 X MMR and the injected doses of mRNAs or MO per cell are as follows: *GFP-EFHC2* and *GFP-EFHC2* R133H (78 pg), *GFP-IFT56* and *GFP-IFT80* (100 pg)^134^, membraneRFP(50 pg), *GLG1-FLAG* for rescue experiment (700 pg), and *GLG1* MO (30 ng) for the knockdown experiment. Live images were captured at stage 23 or stage 25 with LSM700 inverted confocal microscope (Carl Zeiss) with a Plan-APOCHROMAT 63×/1.4 oil immersion objective or Nikon eclipse Ti confocal microscope with 60×/1.4 oil immersion objective. Imaging analysis was performed using Fiji. Bonferroni-adjusted *p*-values were calculated in R using the base stats package. Unprocessed, uncompressed raw images are available from Mendeley (doi:10.17632/fnv33t96vn.1).

## SUPPLEMENTAL FIGURES AND LEGENDS

**Figure S1.**
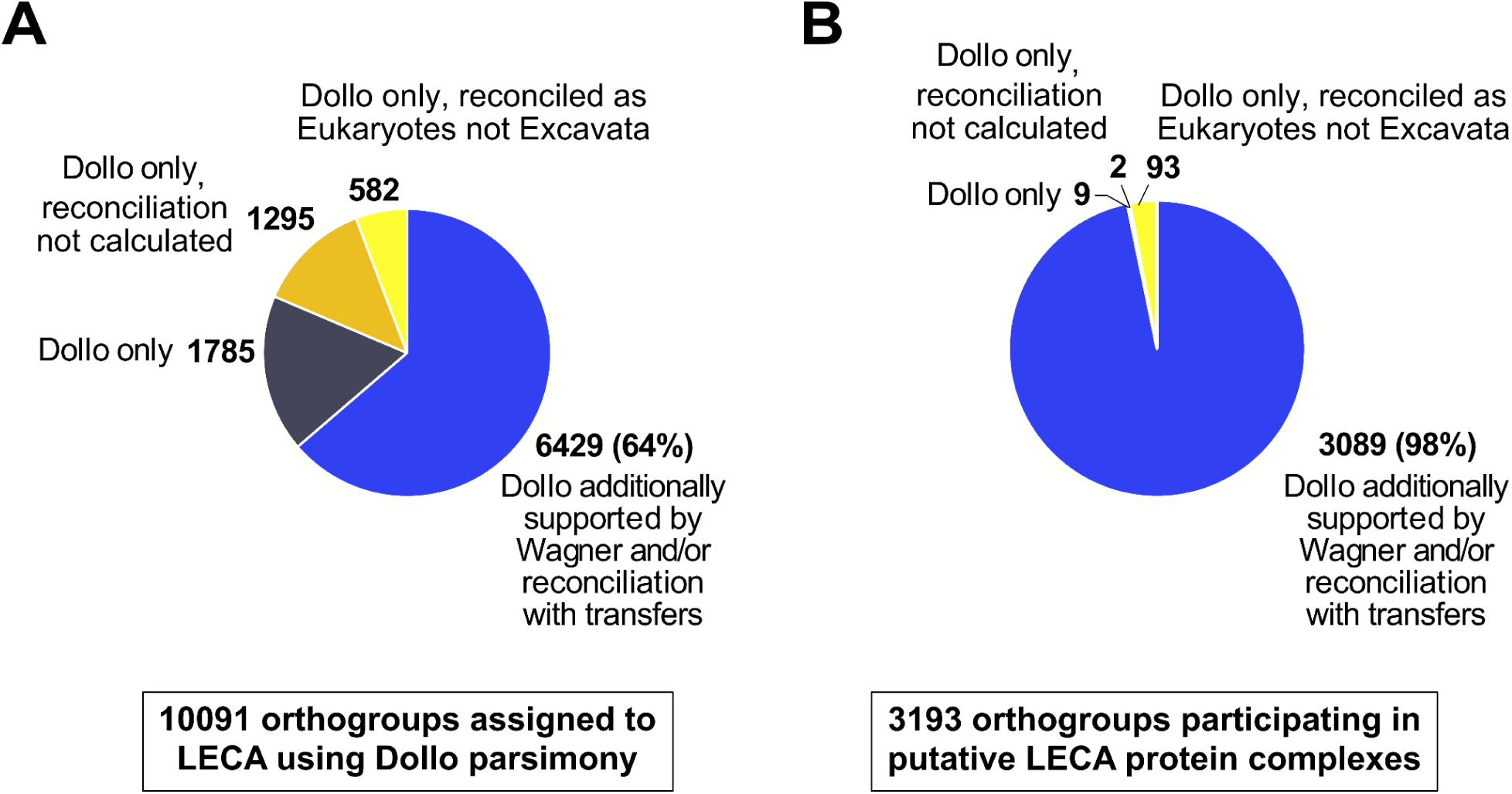
Support for LECA OGs from Wagner parsimony and phylogenetic reconciliation. All OGs presented are supported as present in LECA by at least Dollo parsimony. We further looked for agreement among multiple independent approaches as a robustness test; pathway and gene-level interpretations in the paper are restricted to the multiply-supported subset, with detailed gene-level results provided in **Table S1. (A)** 64% of all LECA OGs supported by Dollo Parsimony are also supported by Wagner or phylogenetic reconciliation with transfers. **(B)** 98% of the core set of 3,193 OGs in the LECA interactome are also supported by Wagner or phylogenetic reconciliation with transfers. For both plots: blue, LECA OGs also supported by Wagner and/or phylogenetic reconciliation with transfers; dark gray, LECA OGs only supported by Dollo parsimony; orange, LECA OGs supported by Dollo parsimony for which phylogenetic reconciliation was not calculated; yellow, LECA OGs supported by Dollo and reconciled as Eukaryotes not Excavata.

**Figure S2.**
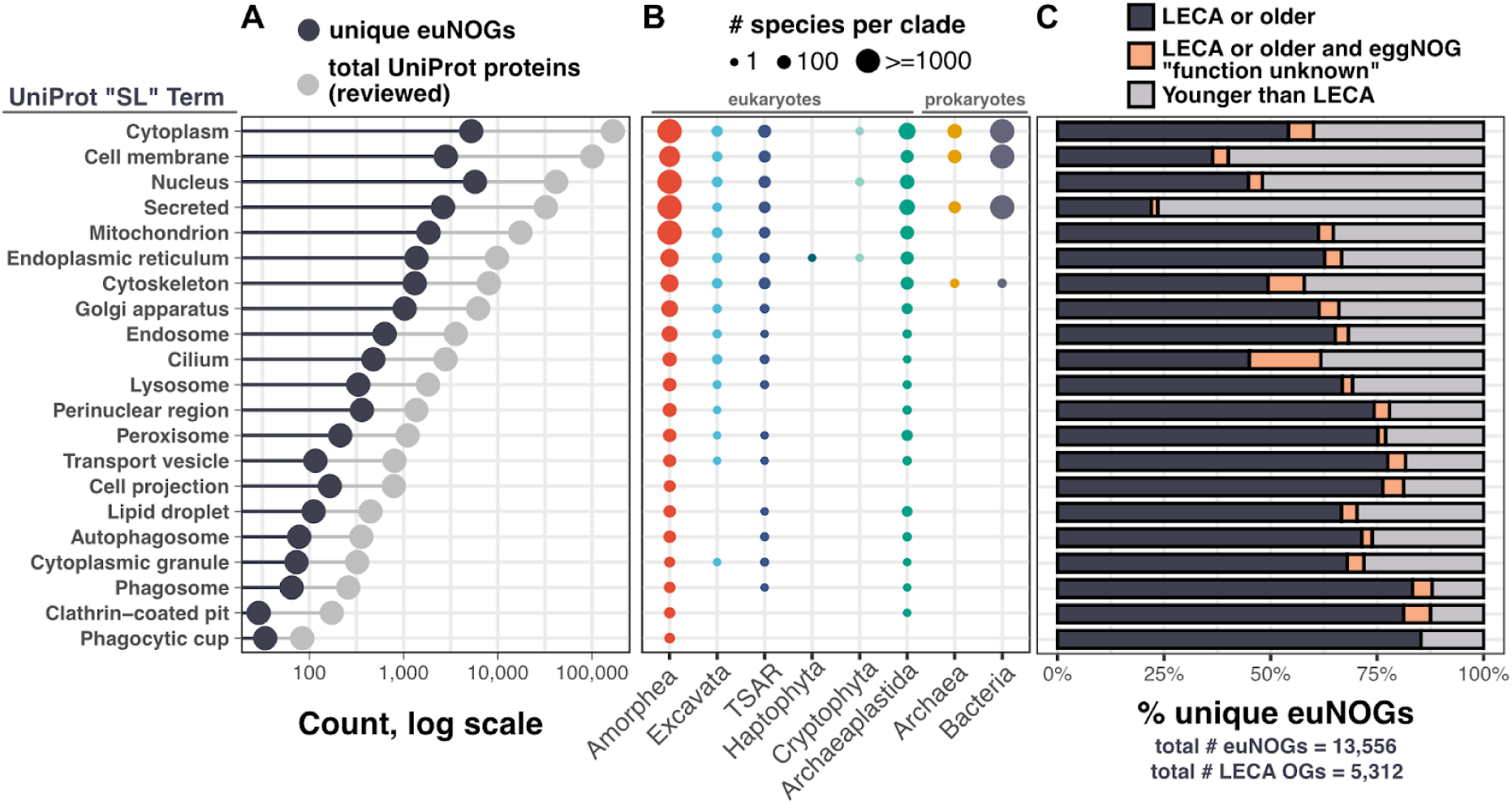
Phylogenetic analysis of the reviewed UniProt database by subcellular localization. Limitations to available annotations are evident in an analysis of the UniProt protein database across species, where reviewed proteins have assigned subcellular localizations likely present in the last eukaryotic common ancestor. **(A)** Light gray, total number of reviewed UniProt proteins by UniProt SL term; dark gray, total number of unique eukaryotic OGs assigned to UniProt proteins by UniProt SL term. **(B)** Phylogenetic representation of the proteins sourced from UniProt by UniProt SL term. **(C)** The percentage of eukaryotic orthologous groups (euNOGs) that trace back to LECA by UniProt SL term.

**Figure S3.**
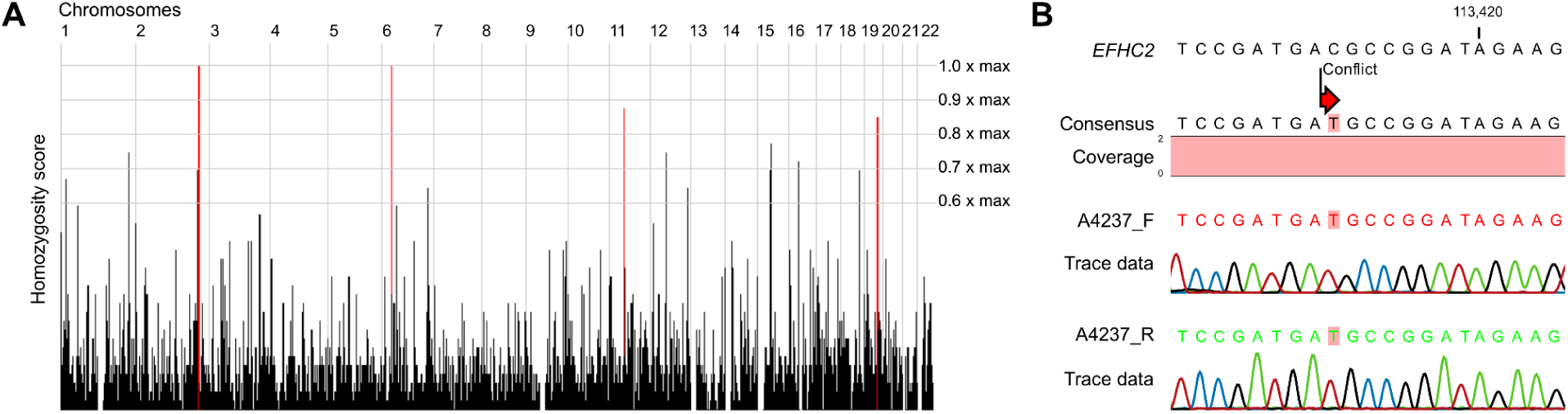
Homozygosity mapping and verification of EFHC2 mutation in individual A4237-22. **(A)** Homozygosity mapping depicts a homozygosity of 4.4 Mb and confirms the reported non-consanguinity of the parents. **(B)** Chromatograms obtained by direct sequencing of PCR products reveal a homozygous substitution of C for T in exon 4 of the *EFHC2* gene in A4237-22.

**Figure S4.**
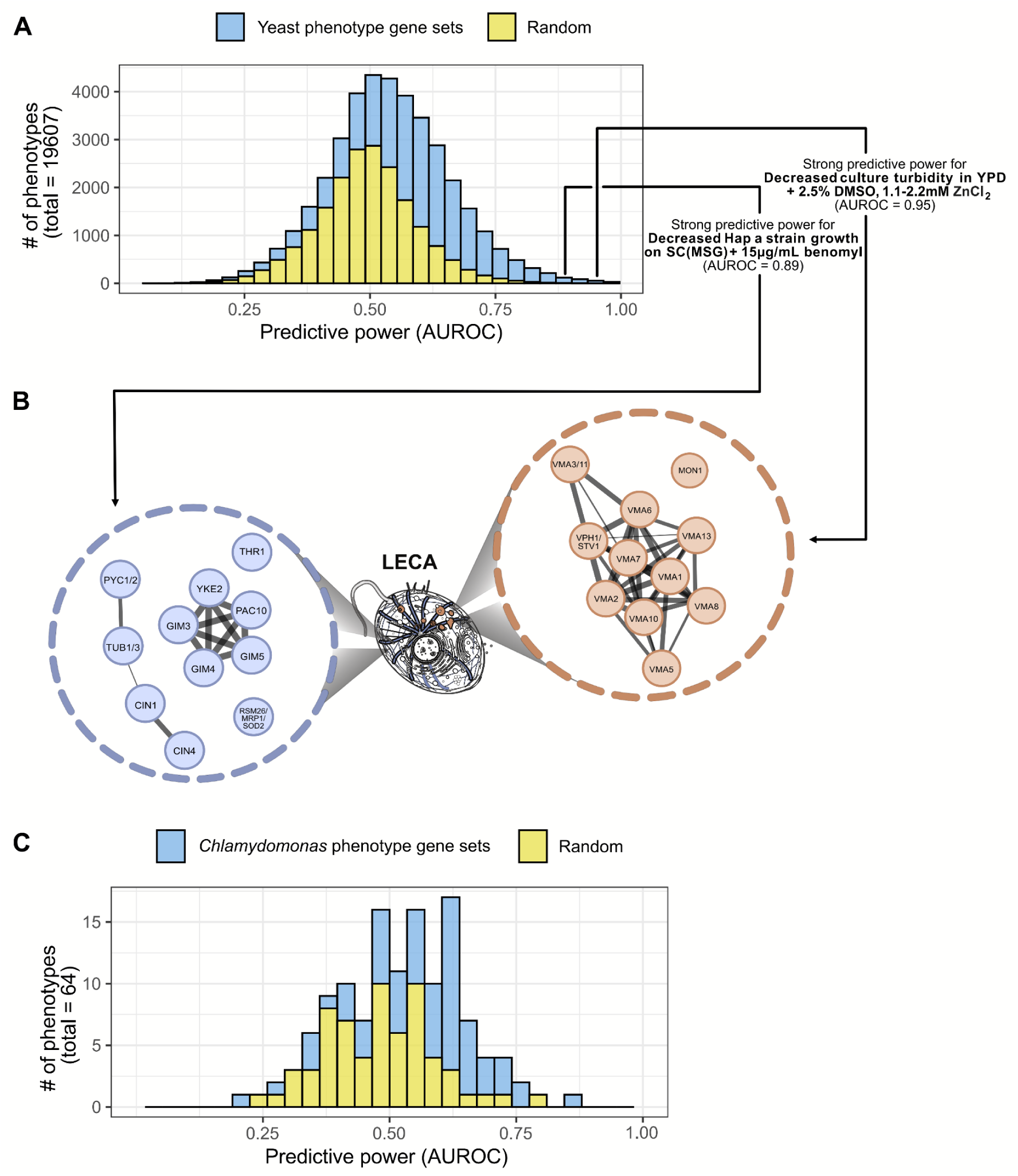
Guilt-by-association in the LECA interactome predicts loss-of-function phenotypes in mutant (A,B) yeast and (C) *Chlamydomonas*. Performance in panels **A** and **C** was quantified using the area under receiver operating characteristic curves (AUROC) for leave-one-out cross-validated predictions of known phenotype-linked genes (light blue) versus random associations (yellow), calculated as for Figure 7A and with predictive performance being roughly comparable to the prediction of the human disease gene sets in Figure 7A. Panel **(B)** shows the relevant LECA PPI networks for genes associated with the two labeled yeast phenotypes.

**Figure S5.**
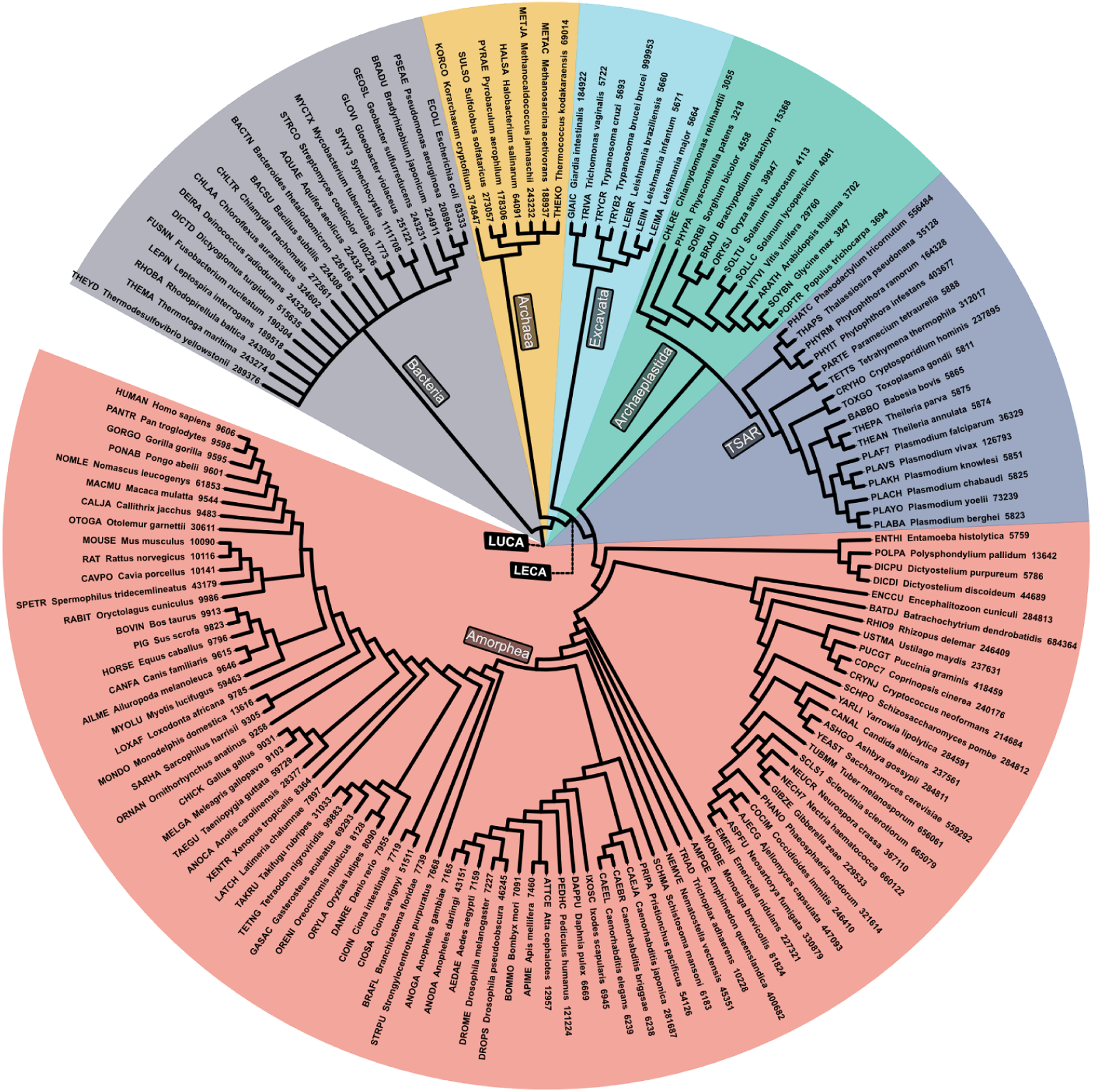
Reference species tree illustration generated by the Interactive Tree of Life for most of the Quest for Orthologs benchmark species (147/156) used in the Dollo and Wagner parsimony analyses. Branch lengths are not to scale. Major supergroups are highlighted across the tree. Prokaryotic groups include Bacteria (gray) and Archaea (yellow). Eukaryotic groups include Excavata (light blue), Archaeplastida (green), TSAR (purple), and Amorphea (red).

**Figure S6.**
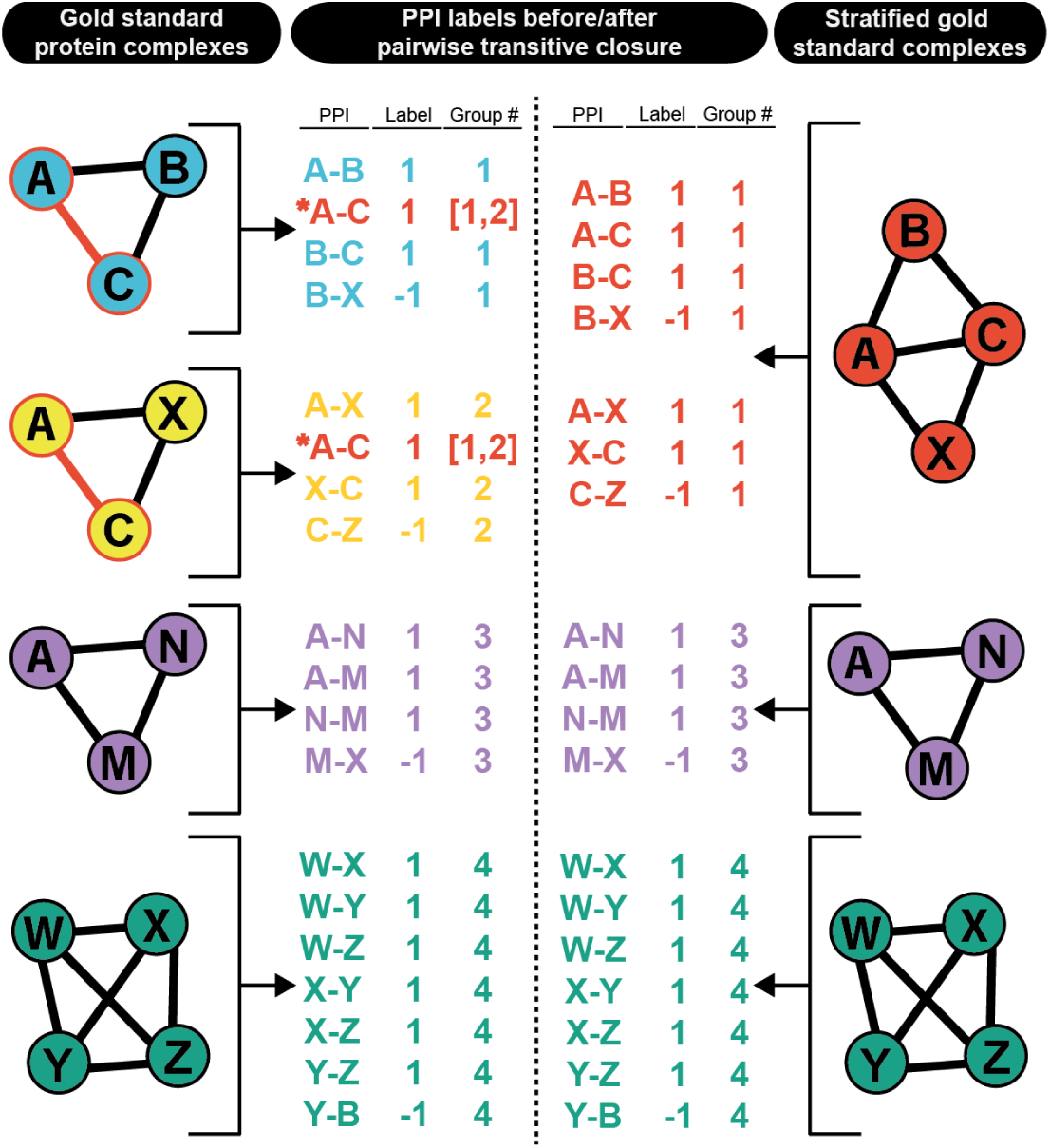
**Illustration of transitive closure for grouping gold standard protein complexes into supergroups.**

**Figure S7.**
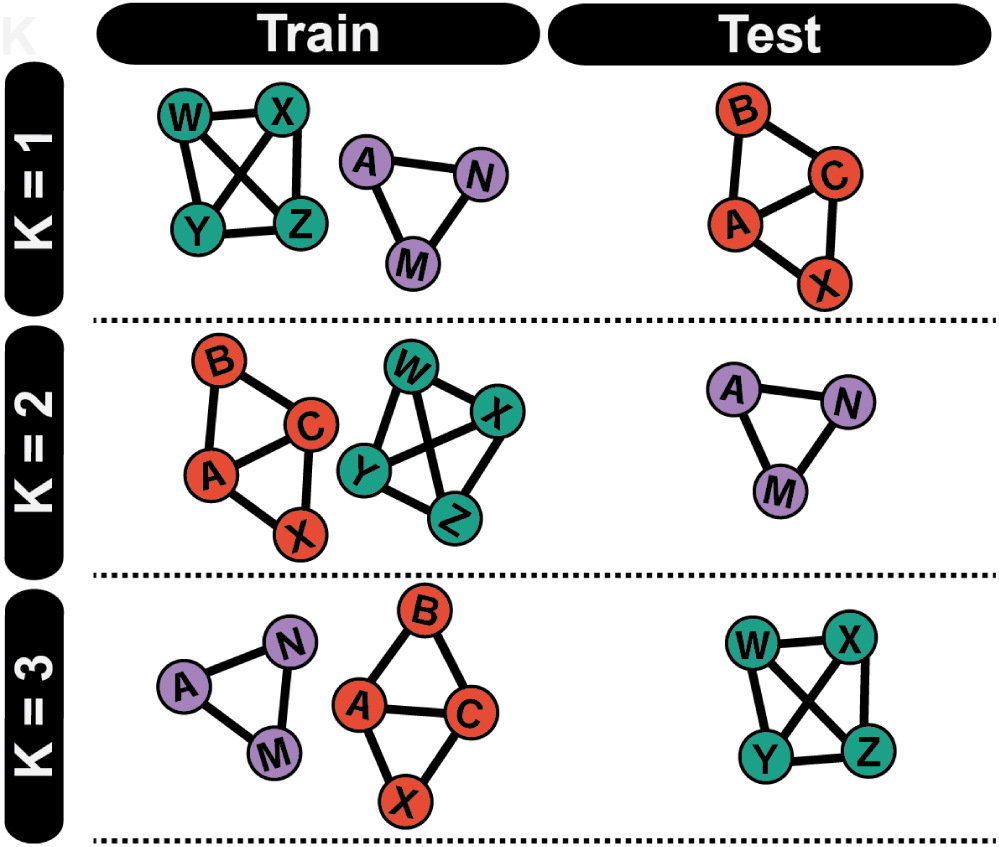
Illustration of group-based k-fold (in this example, k=3) cross-validation for protein-protein interactions.

**Figure S8.**
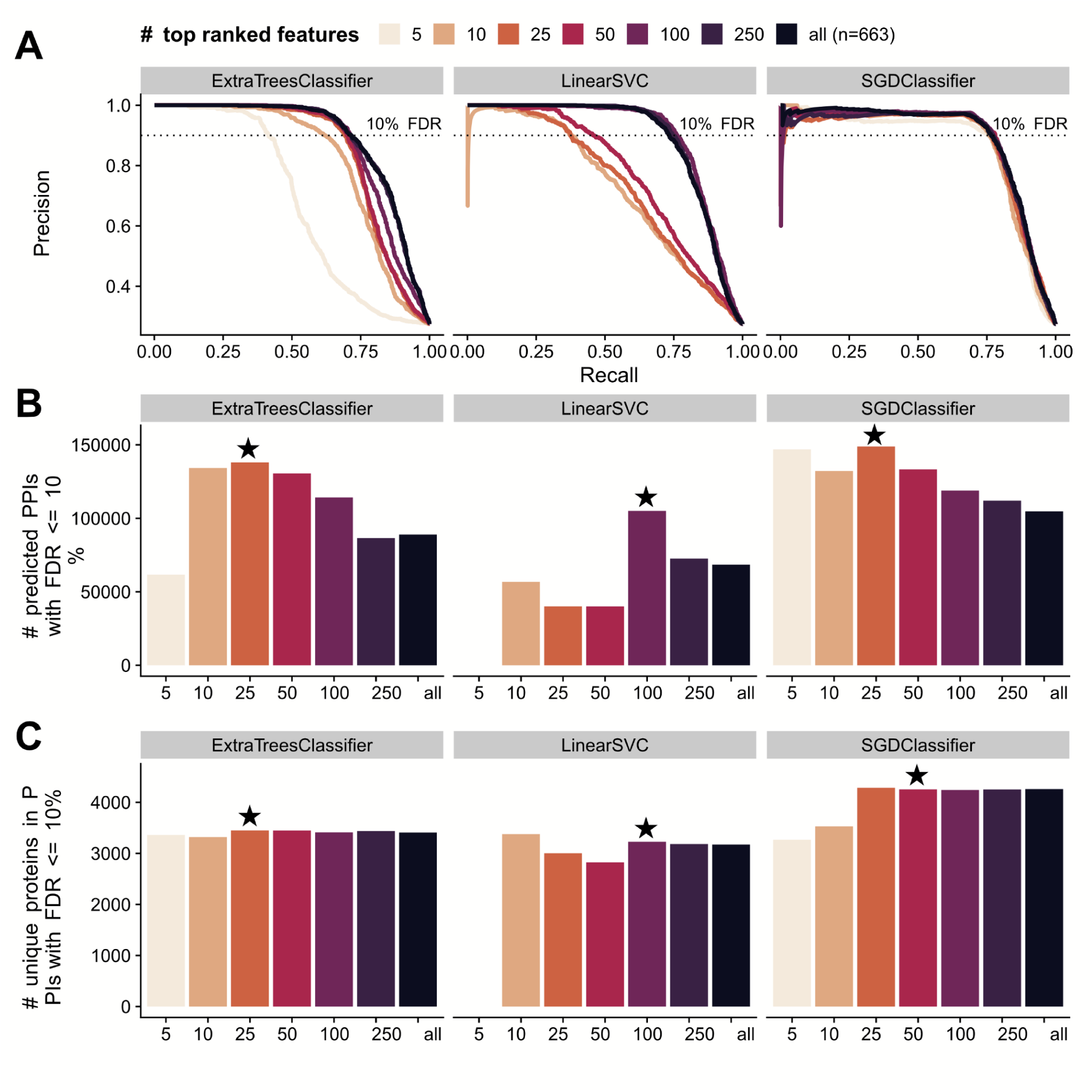
Model selection and optimization. (A) Precision-recall curves for three different algorithms, varying the number of “top” most important features used as input (duplicate panel to Figure 3B). Feature importance is defined per algorithm, ranked by either the absolute value of coefficients for linear models (LinearSVC, SGDClassifier) or the Gini index (ExtraTreesClassifier). (B) The number of pairwise protein-protein interactions (PPIs) within a 10% FDR threshold for each model. Black stars (★) denote the final models used as input to a community detection algorithm to define protein complexes. (C) The number of unique proteins that have at least one interaction scored within a 10% FDR threshold for each model.

### Supplemental Tables

**Table S1.** 10,091 orthogroups estimated to have been present in LECA with associated support according to Dollo parsimony, Wagner parsimony, and phylogenetic reconciliation. 12MB file, available on Zenodo repository. (Note that some cells exceed MS Excel text size limits.)

**Table S2.** 3,193 LECA orthogroups organized hierarchically into 2,013 protein assemblies. 1.3 MB file, available on Zenodo repository.

**Table S3.**
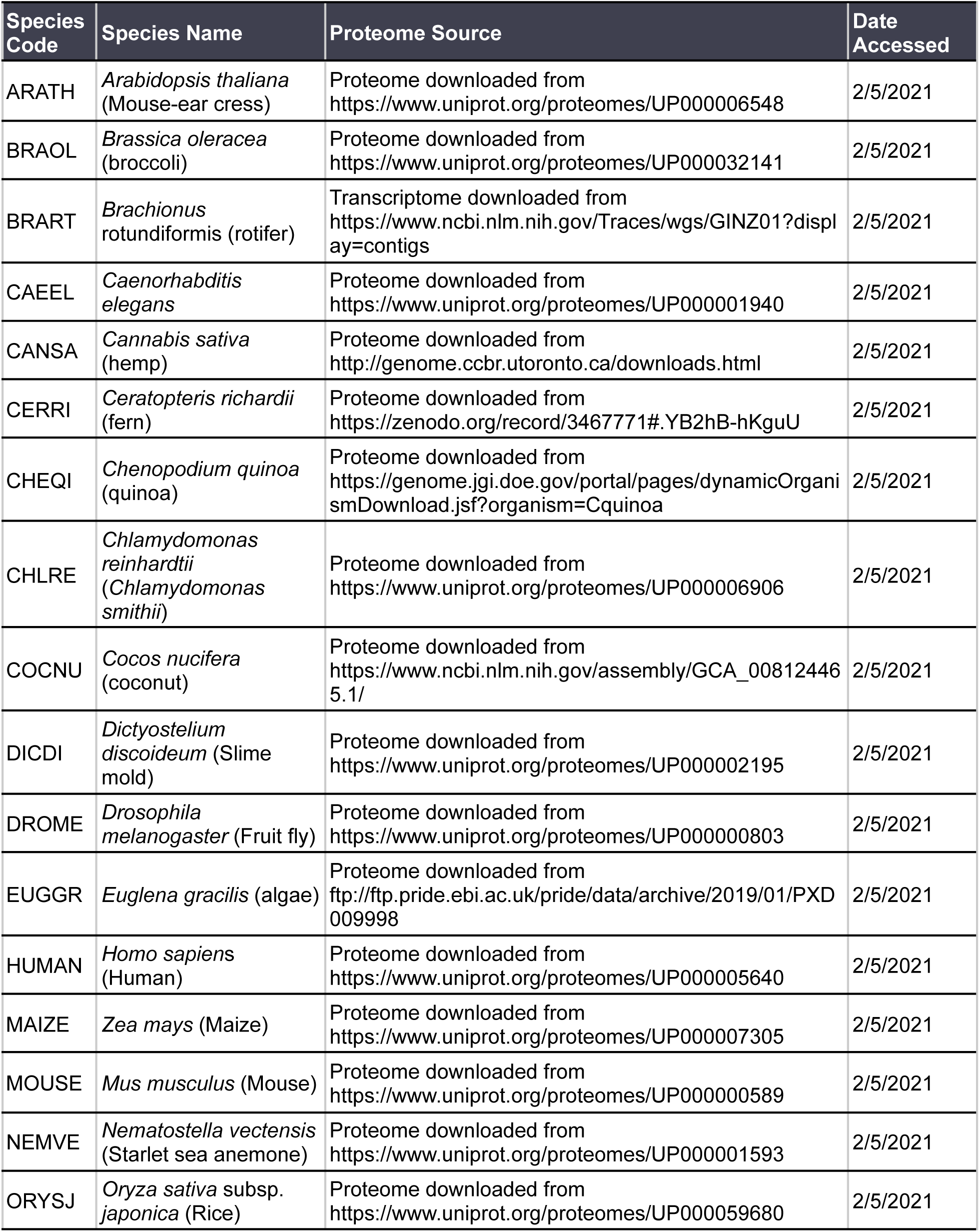
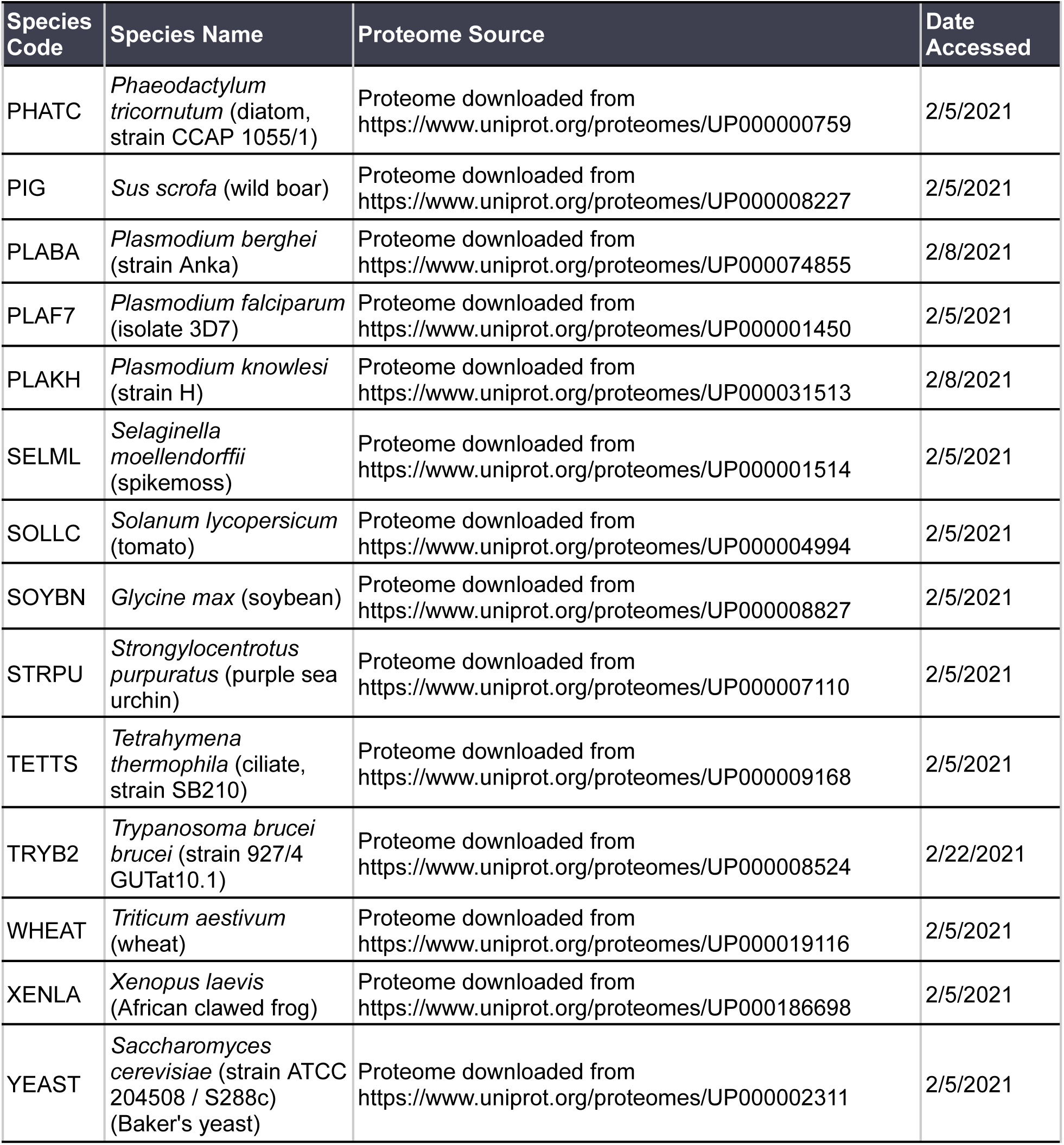
Reference proteomes sourced for co-fractionation mass spectrometry data processing.

**Table S4.**
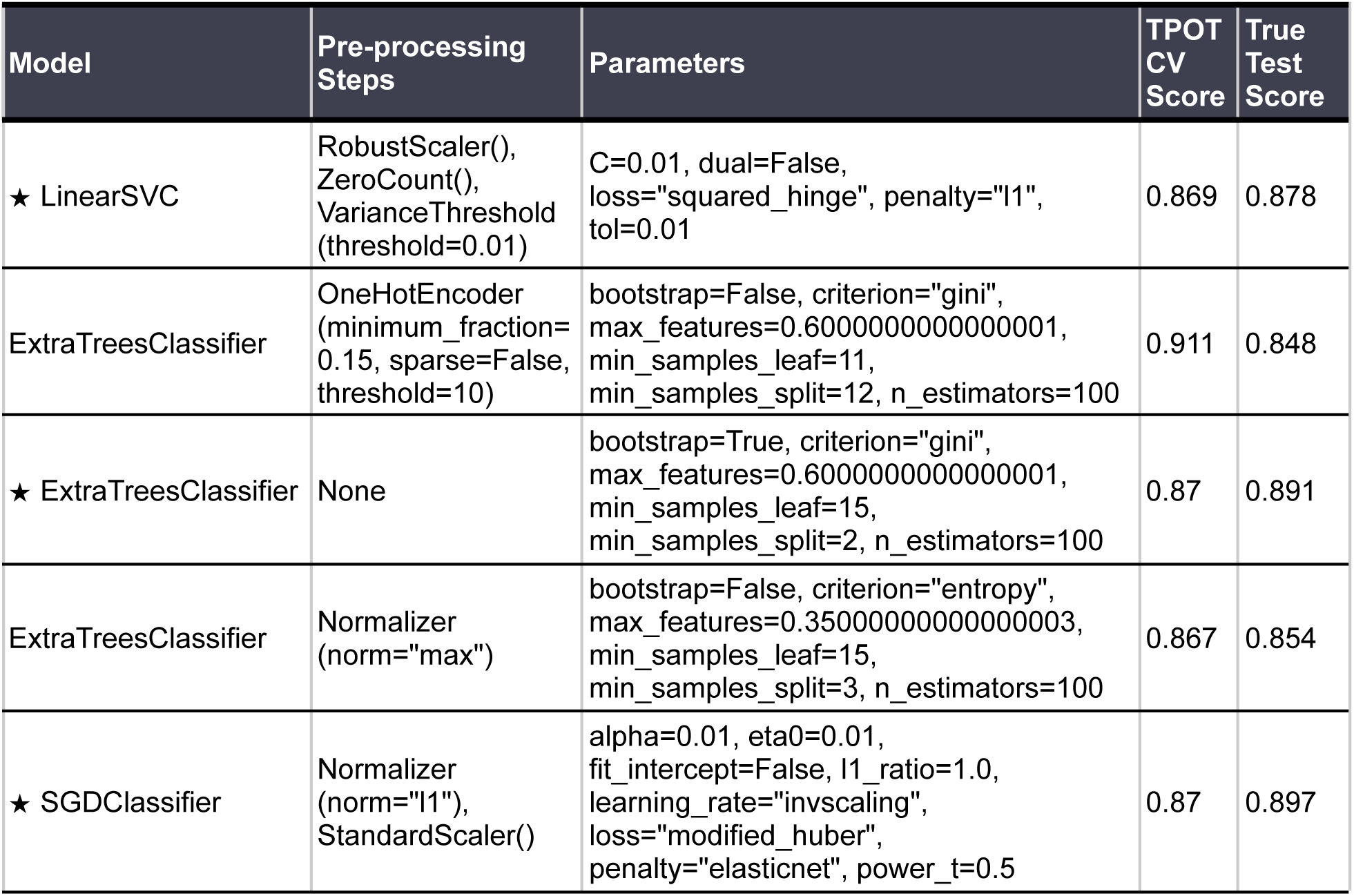
Summary of the top scoring algorithms, parameters and pre-processing steps found by TPOT. . Models marked with a star (★) were selected for further evaluation.

| Model | Pre-processing Steps | Parameters | TPOT CV Score | True Test Score |
| --- | --- | --- | --- | --- |
| ★ LinearSVC | RobustScaler(),<br>ZeroCount(),<br>VarianceThreshold<br>(threshold=0.01) | C=0.01, dual=False,<br>loss="squared_hinge", penalty="l1",<br>tol=0.01 | 0.869 | 0.878 |
| ExtraTreesClassifier | OneHotEncoder<br>(minimum_fraction=<br>0.15, sparse=False,<br>threshold=10) | bootstrap=False, criterion="gini",<br>max_features=0.6000000000000001,<br>min_samples_leaf=11,<br>min_samples_split=12, n_estimators=100 | 0.911 | 0.848 |
| ★ ExtraTreesClassifier | None | bootstrap=True, criterion="gini",<br>max_features=0.6000000000000001,<br>min_samples_leaf=15,<br>min_samples_split=2, n_estimators=100 | 0.87 | 0.891 |
| ExtraTreesClassifier | Normalizer<br>(norm="max") | bootstrap=False, criterion="entropy",<br>max_features=0.35000000000000003,<br>min_samples_leaf=15,<br>min_samples_split=3, n_estimators=100 | 0.867 | 0.854 |
| ★ SGDClassifier | Normalizer<br>(norm="l1"),<br>StandardScaler() | alpha=0.01, eta0=0.01,<br>fit_intercept=False, l1_ratio=1.0,<br>learning_rate="invscaling",<br>loss="modified_huber",<br>penalty="elasticnet", power_t=0.5 | 0.87 | 0.897 |

